# Investigating the conformational dynamics of Zika virus NS4B protein

**DOI:** 10.1101/2021.12.08.471718

**Authors:** Taniya Bhardwaj, Prateek Kumar, Rajanish Giri

## Abstract

Zika virus (ZIKV) NS4B protein is a membranotropic protein having multifunctional roles such as evasion of host-immune system, and induction of host membrane rearrangement for viral genome replication and processing of polyprotein. Despite its versatile functioning, its topology and dynamics are not entirely understood. Presently, there is no NMR or X-ray crystallography-based structure available for any flaviviral NS4B protein. Therefore, in this study, we have investigated the structural dynamics of Zika Virus NS4B protein through 3D structure models using molecular dynamics (MD) simulations approach and experiments. Subsequently, we employed a reductionist approach to understand the dynamics of ZIKV NS4B protein. For this, we studied its N-terminal (residues 1-38), C-terminal (residues 194-251), and cytosolic (residues 131-169) regions in isolation. Further, we have performed experiments to prove the maximum dynamics in its cytosolic region. Using a combination of computational tools and circular dichroism (CD) spectroscopy, we validate the cytosolic region as an intrinsically disordered protein region (IDR). The microsecond-long all atoms molecular dynamics and replica-exchange MD simulations complement the experimental observations. We have also analysed its behaviour under the influence of differently charged liposomes and macromolecular crowding agents which may have significance on its overall dynamics. Lastly, we have proposed a ZIKV NS4B protein model illustrating its putative topology consisting of various membrane-spanning and non-membranous regions.

**Highlights:**

1. Microsecond simulations of NS4B N-terminus and cytosolic regions exposed their dynamic nature.
2. C-terminal region remains intact in presence lipid bilayer during 1 μs simulations.
3. Spectroscopic results also reveal the cytosolic region as an intrinsically disordered region.
4. Cytosolic region exhibits gain-of-structure property under helix inducing solvents.

## Introduction

Flaviviral non-structural 4B protein (NS4B) is a membranotropic protein responsible for a plethora of cellular and molecular functions. The multifunctionality of this protein encompasses the virus-specific roles such as induction of membrane rearrangement for viral genome replication inside host cells [1]. It plays a pivotal role in evading host immune responses by downregulating the key interferon pathways centrally engaged in antiviral response by the host [2–5]. It also functions in obstructing the major components of RNA interference pathways such as Dicer, Drosha, Ago1, and Ago2 proteins [5]. Specifically, Zika virus (ZIKV) NS4B protein has been reported to impede neurogenesis and induce autophagy via inhibiting the Akt/mTOR signaling pathway [6]. Thus, it is suggested to appear as a chameleon with substantial anti-host functions, yet, it remains one of the understudied proteins of flaviviruses. To this end, the membrane topology of NS4B protein of only Dengue virus (DENV) and West Nile Virus (WNV) has been established experimentally [7–9], but for ZIKV NS4B, not enough evidences are available.

According to the topology model of flaviviral NS4B protein, it comprises of transmembrane regions with five domains spanning the membrane of the endoplasmic reticulum (ER) in hosts [7,8]. The model suggests the presence of two predicted and three proven transmembrane domains (TMDs): pTMD1, pTMD2, TMD3, TMD4, and TMD5 [7]. In DENV2, a loop of ~35 residues connecting TMD3 to TMD4 occupies the cytoplasmic region, which is reported to indulge in dimer formation [9]. This region also functions in viral replication and acts as a cofactor for the helicase activity of DENV NS3 protein [10,11]. Furthermore, it has been shown to dissociate the newly formed single-stranded viral RNA from NS3 helicase and enhance its unwinding property [10,11]. Despite of its multifunctionality, multiple aspects such as topology, folding, structural conformations, and interactions are yet to be studied. Therefore, we have aimed to understand the conformation of ZIKV NS4B and its regions forming rigid and flexible conformations within the protein.

Careful analysis and comparison of proteins from representative organisms of all three kingdoms of life exposed the existence of intrinsically disordered regions (IDR)-containing proteins in them [12]. Another benchmark study correlated the connection between the disordered content and proteome complexity of thousands of species from three domains of life and viruses. Particularly, viruses are highly enriched in intrinsic disorder and have the most extensive variation range for accommodating IDRs in their proteomes [13]. Viruses heavily use IDRs in proteins to deal with their hostile habitats, also reflecting the ongoing evolution. Several viruses have been studied for the presence of IDRs which have been found associated with their pathogenicity [13]. To name a few, Rotavirus, Chikungunya virus, and members of *Flaviviridae* family such as Hepatitis C virus, Japanese Encephalitis virus (JEV), ZIKV, and DENV, all contain IDRs in their proteins [14–19]. In flaviviruses specifically, the capsid involved in the viral particle maturation is the most disordered protein [17,19,20]. Other proteins like PrM and NS1 also contain a considerable amount of disorder. Due to membrane-spanning regions in flaviviral NS4B protein, it is considered a structured protein where the previously evaluated disorder propensities of ZIKV, DENV, and JEV NS4B proteins are 6.4, 14.5, and 6.66, respectively [17,19,20].

This manuscript aims to understand the dynamics of the full-length ZIKV NS4B protein (NS4B-FL) along with a specific experimental focus on its cytosolic region (NS4B-CR; residues 131-169). According to the findings, the N-terminus (NS4B-NT; residues 1-38) is one of the highly flexible regions while C-terminus (NS4B-CT; residues 194-251) residues have tendency to form disordered conformations in absence of membrane. The cytosolic region connecting the transmembrane domains 3 and 4 residing in the host cytoplasm is another extremely dynamic segment forming an intrinsically unstructured protein region. It also has a gain-of-structure property and can adopt secondary structure while interacting with its physiological partner.

## Experimental Procedures

### Materials

Chemically synthesized ZIKV NS4B cytosolic region (NS4B-CR, 131-169 residues of NS4B-FL protein; ZIKV strain UniProt ID: A0A024B7W1) (NH2-AARAAQKRTAAGIMKNPVVDGIVVTDIDMTIDPQVEKK-COOH) with 96% purity was purchased from Thermo scientific USA and dissolved in buffer consisting of 20 mM sodium phosphate buffer (pH 7.4). Chemicals including Sodium Dodecyl Sulfate, 2,2,2-Trifluoroethanol, Trimethylamine *N*-oxide, and Poly Ethylene Glycol 8000 were obtained from Sigma Aldrich, St. Louis, USA. Ficoll-70 was purchased from Cytiva, Marlborough, MA, USA. Liposomes were purchased from Avanti Polar Lipids, Inc., Alabaster, AL, USA.

### Computational methods

#### Multiple sequence alignment

To map the residues forming the cytoplasmic region in ZIKV NS4B protein, multiple sequence alignment of full-length NS4B proteins of different flaviviruses – DENV (all four serotypes), JEV, WNV, YFV, and KUJV was done using Clustal Omega [21] with default parameters.

#### Secondary structure and transmembrane prediction

So far, no crystal structure or NMR structure has been determined for any flaviviral NS4B protein. Therefore, we employed a combination of two different predictors, TMPred [22] and TMHMM [23], to predict the presence of transmembrane regions in ZIKV NS4B protein. To predict the secondary structure of NS4B-CR in isolation, PSIPRED was used [24]. These web predictors use highly optimized and least biased algorithms to give an overall secondary structure of the input sequence.

#### 3D Model building

To gain insight in three-dimensional space, the 3D models were built to explore the structural dynamics of NS4B-FL (full-length) and NS4B-NT (N-terminal 1-38 residues), NS4B-CR (cytosolic region 131-169 residues), and NS4B-CT (C-terminal 194-251 residues) regions in isolation. We have utilized I-TASSER [25] and recently introduced artificial intelligence-based AlphaFold2 software for model building [26]. In I-TASSER, based on high confidence score (C-score) the best model is chosen. Similarly, the AlphaFold2 has produced five high quality models, out of which the first ranked model is chosen for further molecular dynamics study.

#### Molecular dynamics (MD) simulations

For all simulations, in the Desmond simulation package, embedded in the Schrodinger suite of drug discovery, we have employed OPLS 2005 forcefield to investigate the dynamics of NS4B-FL, NS4B-NT, NS4B-CT and NS4B-CR in isolation. Multiple simulations of all models were performed to one microsecond (μs). Since flaviviral NS4B is a transmembrane protein, we have performed its simulation in the presence of lipid bilayer. we have placed its membrane-spanning residues (39-59, 79-96, 107-127, and 173-193) in a lipid bilayer having POPC (1-palmitoyl-2-oleoyl-sn-glycero-3-phosphocholine) molecules surrounded by TIP3P water models. Simulations of models of NS4B-NT and NS4B-CR segments in isolation is performed only in absence of lipid bilayer. NS4B-CT model is simulated in presence as well as in absence of POPC membrane. We have used the previously reported methodology of MD simulations of isolated regions like the C-terminal or cytosolic region of SARS-CoV-2 Spike protein [27].

Further, the simulation trajectory of the I-TASSER model of NS4B-CR was subjected to RMSD clustering. For RMSD clustering using the *Desmond trajectory clustering* tool, the microsecond long trajectory is analyzed. Further, the Replica Exchange MD (REMD) simulations are performed for 500 nanoseconds (ns) with previously used protocol in the Desmond simulation package using the last simulated frame of cytosolic region from I-TASSER simulation trajectory (i.e., 1 μs) [27]. Here, we have investigated the conformational dynamics of NS4B-CR with 12 replicas at increasing temperatures, ranging from 300K to 410K at a regular interval of 10K.

#### Prediction of intrinsic disorder propensity

A total of three widely used online predictors were utilized to predict the intrinsic disorder predisposition in ZIKV NS4B-CR. IUPred2A platform (long and short) [28], DISOclust [29], and PONDR (prediction of naturally disordered regions) family such as PONDR-VSL2, -VLXT, and - VL3 were used [30]. One meta-predictor - PONDR-FIT was also employed to enhance the disorder analysis [31]. Disorder propensities from all predictors were used to calculate the mean disorder predisposition in NS4B-CR.

### Experimental methods

#### CD spectroscopy

Far-UV CD spectra of samples were measured in 1.0 mm path length quartz cuvette from 190 nm to 240 nm wavelength using the J-1000 series JASCO (Easton, MD, USA) spectrophotometer system. The bandwidth was set at 0.5 mm, and values of three spectra were averaged to obtain a single spectrum. We obtained the spectra at 25 °C in the presence of 20 mM sodium phosphate buffer, pH 7.4. Buffer baselines were measured to normalize the CD scans of samples. For temperature-dependent far-UV CD scans, spectra were measured at every 10 °C from 30 °C to 90 °C.

#### Preparation of liposomes

Large unilamellar vesicle (LUVs) or liposomes distinctly composed of 1,2-dioleoyl-sn-glycero-3-phosphocholine (DOPC), 1,2-dioleoyl-sn-glycero-3-phospho-L-serine (DOPS), and 1,2-dioleoyl-3-trimethylammonium-propane (DOTAP) were prepared following the thin-film hydration method briefly described in our previous reports [19].

#### Dynamic light scattering

The hydrodynamic radius of prepared LUVs was measured through dynamic light scattering using Zetasizer Nano ZS (Malvern Panalytical Ltd., Malvern, UK). The instrument is equipped with a laser (633 nm) set at an angle of 173°. Equilibration time of 60 seconds and 10 seconds of run time was chosen for measurements. Three measurements were recorded (chosen ten times run) for each sample. Prepared LUVs were diluted in water (5:100 ratio of LUVs and water) before taking the measurements.

## Results

### 1. Prediction of secondary structure of ZIKV NS4B

The topology of flaviviral NS4B protein is not yet completely understood. Miller *et al*., have characterized DENV NS4B protein and suggested a model for its topology as a membrane protein residing in the endoplasmic reticulum membrane [7]. Here, we have predicted the putative topology of ZIKV NS4B-FL using the topmost available predictors TMHMM and TMPred (**Figure S1**). TMHMM has predicted three transmembrane regions: residues 39-54, 74-81, and 108-124 in NS4B protein which satisfies the default threshold (probability of 0.8 and above). Two transmembrane regions are also present near the C-terminus: residues 174-190, and 230-239 which shows slightly lower values than the threshold (probability values between 0.7-0.8) (**Figure S1A**). TMPred results also indicate the presence of similar five transmembrane regions. The residues 39-58, 66-89, 104-122, 170-192, and 226-248 are predicted to form transmembrane helices in the strongly preferred model given by TMPred. In an alternate model of TMPred, only four transmembrane helices from residues 39-55, 65-89, 103-123, and 170-188 are predicted (**Figure S1B**).

### 2. Structural dynamics of NS4B protein and its N & C terminal regions

#### 2.1. MD Simulation of full-length NS4B protein

To investigate the structural dynamics of NS4B-FL protein, we constructed structural models of NS4B-FL using AlphaFold2 and I-TASSER which were simulated for 1 μs in presence of POPC membrane. **Figure 1** depicts both the models of ZIKV NS4B-FL and their simulation setup. The dynamics of both predicted models through different frames at every 100 ns of the simulation trajectory is illustrated in **figure 2**.

**Figure 1:**
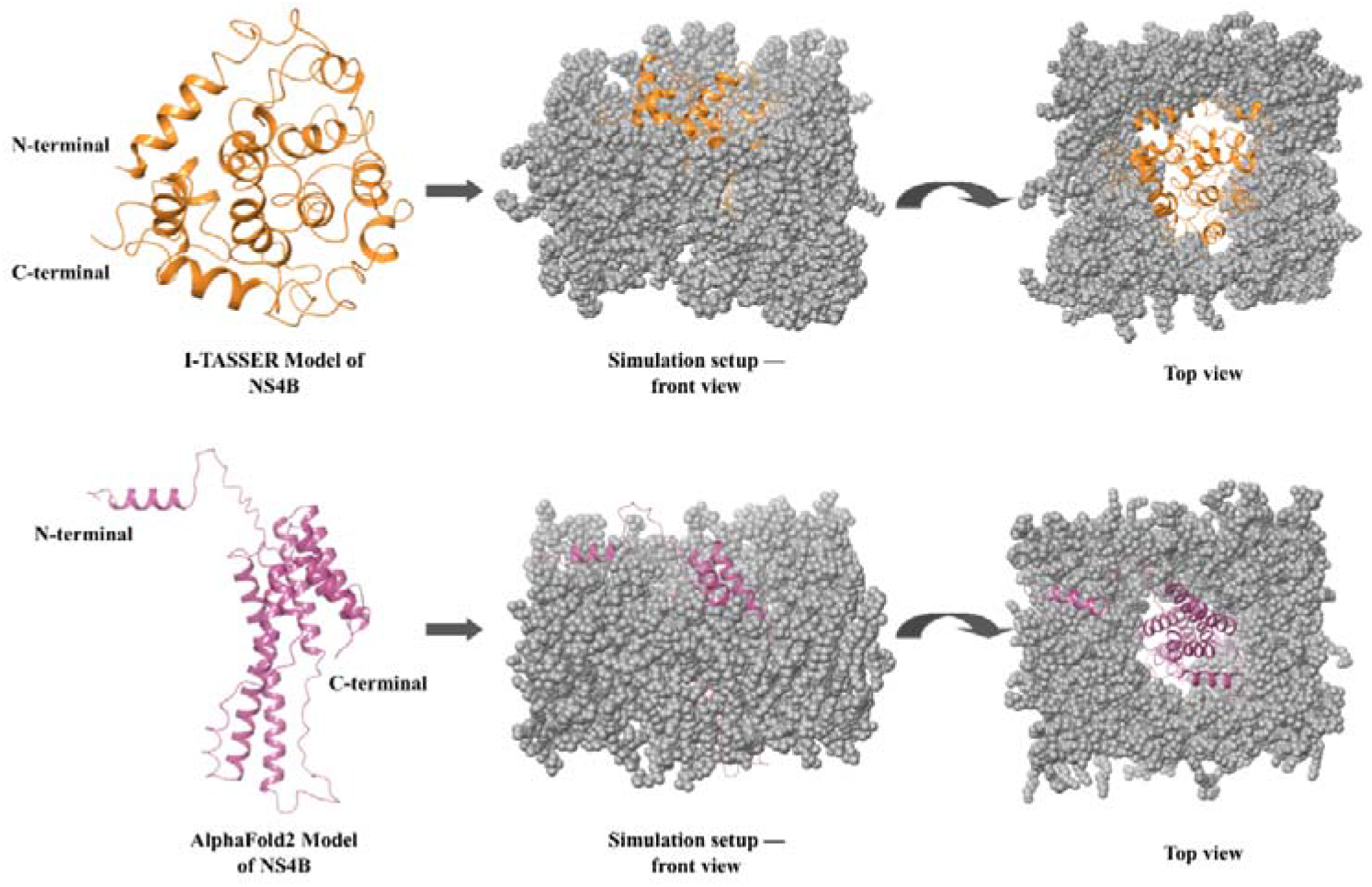
3D structure models of full-length ZIKV NS4B, generated by I-TASSER (orange) and AlphaFold2 (magenta), are simulated for 1 μs in the presence of POPC membrane (grey).

**Figure 2:**
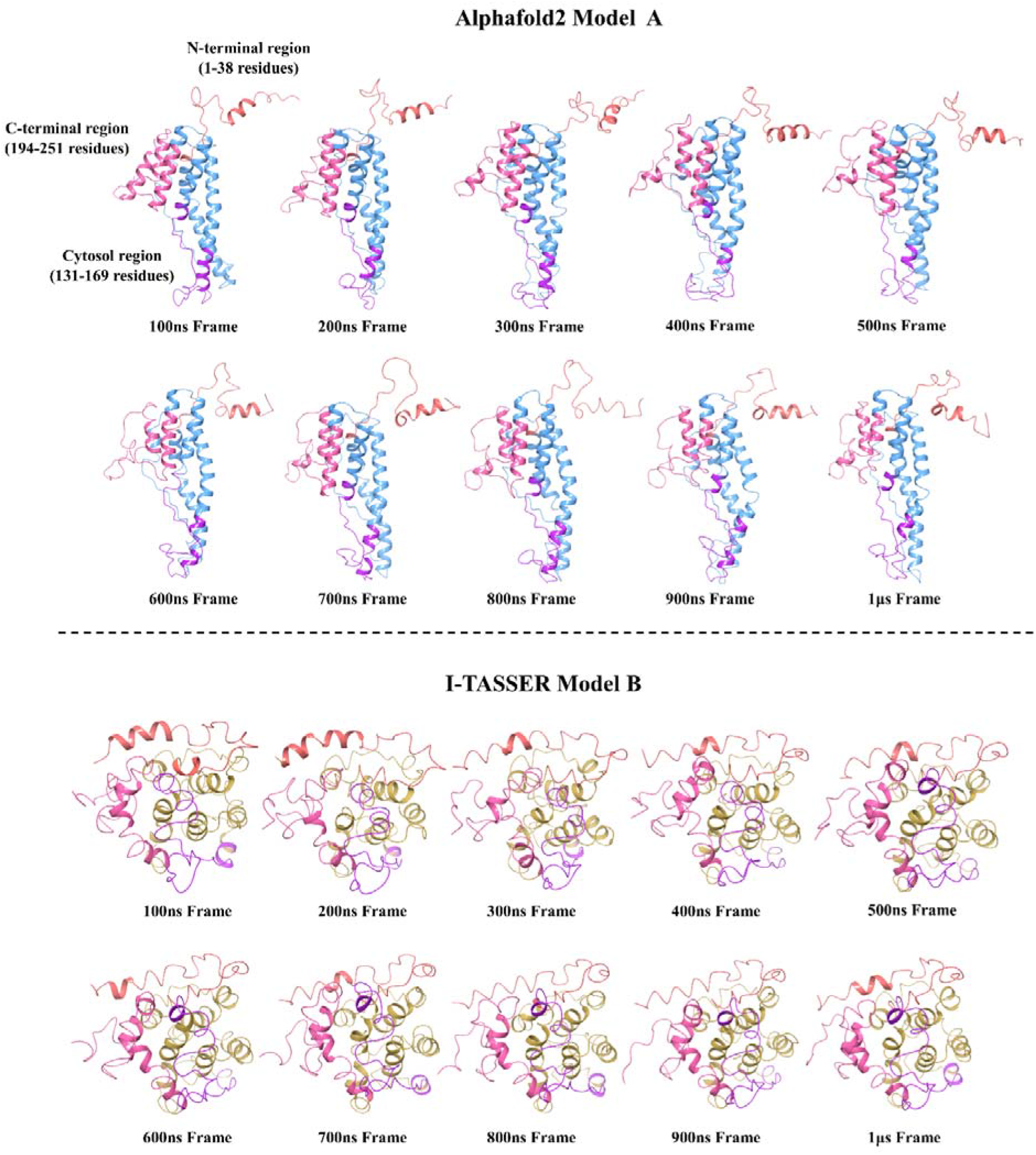
Snapshots from 1 μs long MD simulations trajectory of predicted model A (AlphaFold2; in upper panel; shown in faded azure) and model B (I-TASSER; in lower panel; shown in golden olive) of ZIKV NS4B-FL protein in the presence of POPC membrane. The N-terminal, C-terminal and cytosolic region are represented with faded red, faded salmon, and plum color respectively in both the modeled structures.

Consequently, both NS4B-FL protein models display significant changes in their structure throughout the simulation period. Based on atomic deviation, RMSD, calculation of each frame with reference to the initial frame in the presence of POPC lipids, the average RMSD of simulation trajectory of AlphaFold2 predicted model A is 5.5 Å. For model B built using I-TASSER, the RMSD value is evaluated to be 3.7 Å. The mean fluctuation values (RMSF) of model A’s 1 μs long simulation trajectory varies up to 5 Å for non-transmembrane regions while transmembrane residues have shown least fluctuations up to 2 Å. In comparison, the model B simulation trajectory has shown less fluctuations ranging between 1-4 Å. Graphs corresponding to RMSD and RMSF for NS4B-FL models are depicted in **figures S2A**, **S2B**, **S3A**, and **S3B**. In both the models, most of the residues of N-terminal (depicted with orange color in **figure 2**), and C-terminal (depicted with pink color in **figure 2**) have gained a disordered conformation. Cytosolic region which is shown in purple color also became unstructured during the course of 1 μs. Particularly in model A, residues (39-59, 79-96, 107-127, and 173-193) which were placed inside the POPC bilayer display relatively less fluctuations (refer to RMSF plots in **figure S2B**). After tertiary structure analysis upon visualization, the secondary structure analysis is consistent with the visualized 3D structure. Overall, the secondary structure element is higher (~47%) in model A than the model B (~28%) and consist of no beta strands. Plots in **figures S2C**, **S2D**, **S3C** and **S3D** summarizes the secondary structure analysis through the timeline for models A and B in supplementary information. In **supplementary movie 1** and **2**, trajectories of simulations of respective models A and B are shown.

#### 2.2. MD simulation of NS4B N-terminal (NS4B-NT)

Next, we studied the dynamics of N-terminal region of ZIKV NS4B protein in isolation. The AlphaFold2 model of NS4B-NT consist of nearly half of the residues (1-21) forming a helical region and other half of the residues in unstructured conformation. After the simulation of 1 μs, the helical regions gradually turned into the unstructured region, as shown in **figure 3**. However, a small helix formed by residues _14_SHLMG_18_ can be seen in most of the snapshots captured at every 100 ns of trajectory. The **supplementary movie 3** shows the 1 μs trajectory of NS4B-NT simulation.

**Figure 3:**
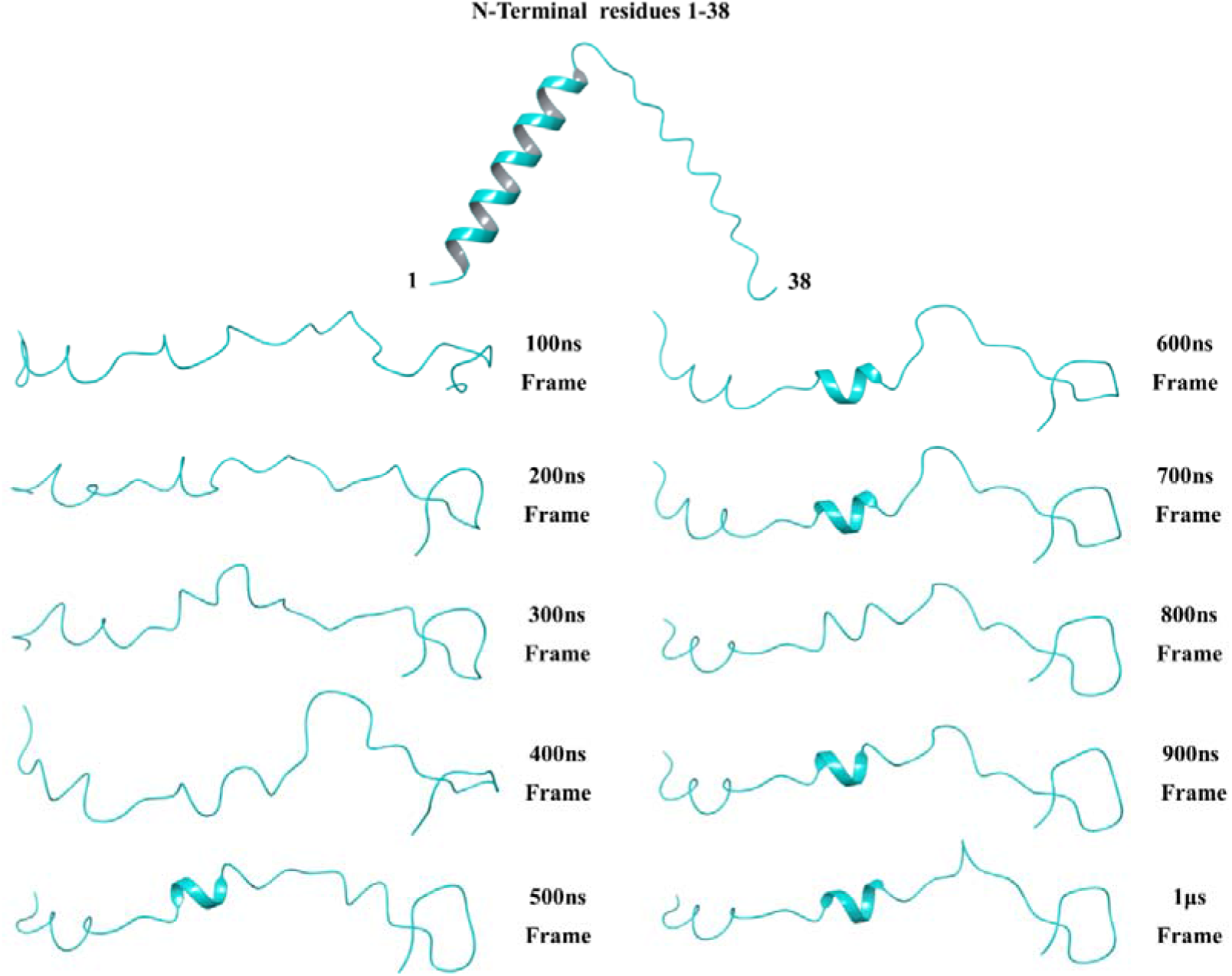
Snapshots of the AlphaFold2 model of ZIKV NS4B N-terminal region (NS4B-NT; residues 1-38; shown in teal color) is simulated for 1 μs. The helix present in the model attains a disordered form during the simulation period.

**Figure S4** further shows the RMSD, RMSF, and secondary structure timeline plots for this simulation. Correspondingly, during the initial course of simulation, the mean deviations are high which stabilizes after 600 ns with an average RMSD of 13 Å (**Figure S4A**). Subsequently, due to high structural transitions, the RMSF values are found to be varying between 2 - 4 Å for all the residues of NS4B-NT (**Figure S4B**). The same has been depicted through the 1 μs timeline and histogram plots for residues in **figures S4C** and **S4D**.

#### 2.3. MD simulation of NS4B C-terminal (NS4B-CT)

Flaviviral NS4B protein is demonstrated to be composed of five membrane-spanning regions out of which one lies in the C-terminus region [7]. Also, the C-terminal region of ZIKV NS4B is predicted to contain a transmembrane region with low probability values (refer to TMHMM and TMPred graphs in supplementary information **figure S1**). Moreover, in DENV system, the C-terminus of NS4B protein has been demonstrated to localize in the ER lumen after cleavage from NS5 by the viral NS2B/NS3 protease [7,9]. Therefore, we have investigated the dynamics of this region in absence as well in presence of POPC membrane.

We have built a 3D model of NS4B-CT in isolation using AlphaFold2 which consists of three helices connected by loops and followed by an unstructured region of 7 residues at extreme C-terminal end. As can be perceived from **figure 4**, in presence of POPC membrane, the structure remains ordered during the simulation period of 1 μs. However, in absence of membrane, the NS4B-CT model gradually loses its helicity and formed an unstructured conformation (**Figure 4**). The flexibility in aqueous environment demonstrate that this region may attain disordered conformation and interact with other macromolecules with its greater radius to perform versatile functions. The statistical analysis of trajectory depicts the changes occurring in 3D structure. Particularly, the NS4B-CT in aqueous environment has experienced high deviations with average RMSD of ~8.7 Å while in hydrophobic environment, the average RMSD is ~2.6 Å (**Figures S5A** and **S6A**). Also, mean residual fluctuations are in good correlation with the snapshots and RMSD values, and clearly describe the structural transitions throughout simulation period. The RMSF values of NS4B-CT in absence and presence of POPC are varying between 2 - 7.5 Å and 0.6 - 2 Å (except the unstructured extreme C-terminal), respectively (**Figures S5B** and **S6B**). Inconsistency or the loss of helicity in hydrophilic environment and intact helices in hydrophobic environment are evident from the secondary structure timelines and histogram plots (**Figures S5C**, **S5D**, **S6C**, and **S6D**). As calculated, less than 20% and more than 60% secondary structure element is observed in absence and presence of POPC in MD simulations, respectively. The trajectories of both the simulations of NS4B-CT are given as **supplementary movies 4** and **5**.

**Figure 4:**
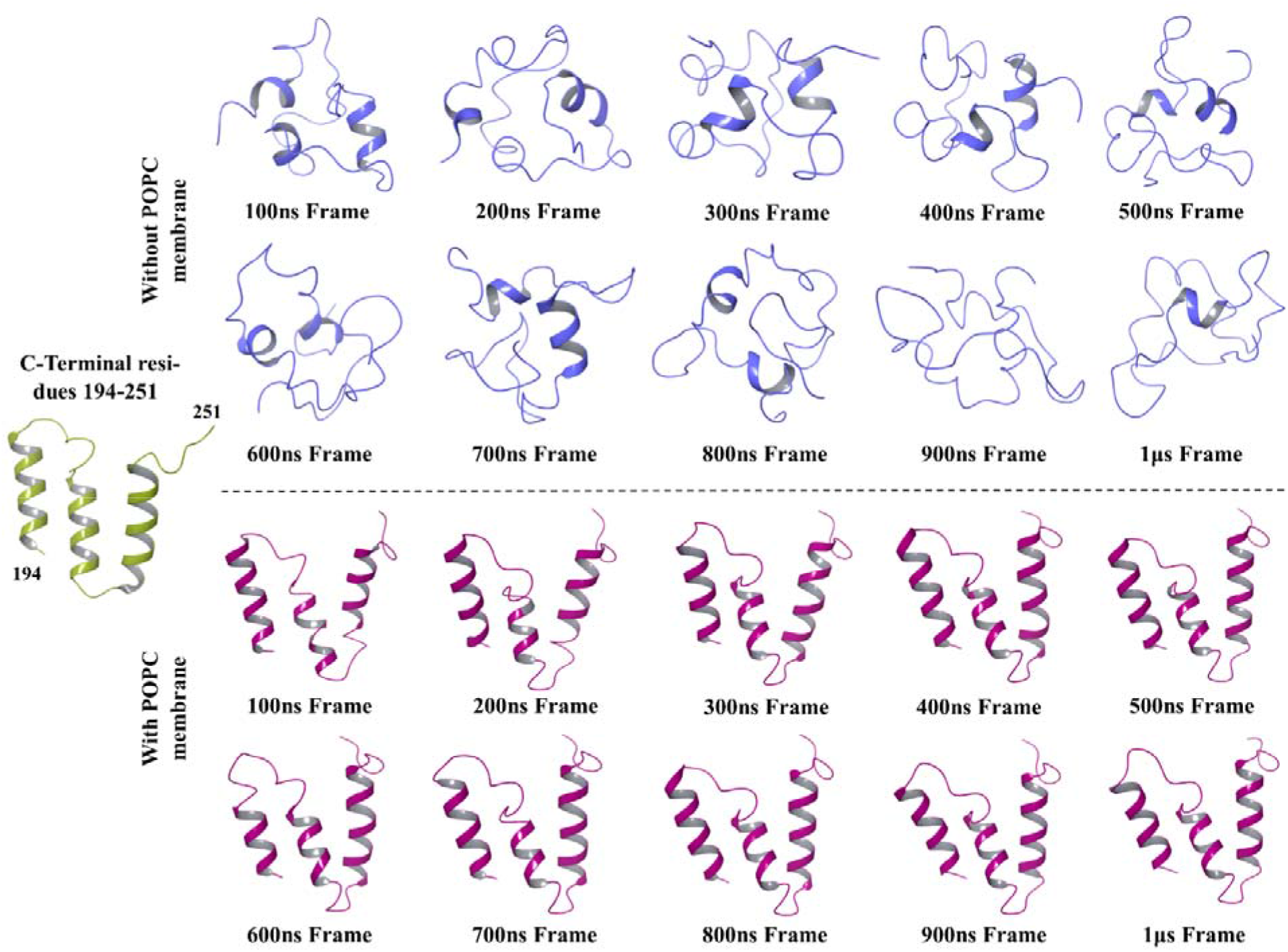
The NS4B-CT model predicted by AlphaFold2 (shown in olive colour) is subjected to 1 μs simulations in absence (cyan colour; in upper panel) and presence (wine colour; in lower panel) of POPC membrane. All snapshots are taken at every 100 ns interval from the trajectory.

### 3. Cytosolic region of NS4B protein

In flaviviral NS4B protein, the residues corresponding to the cytosolic region vary; therefore, we performed a multiple sequence alignment (MSA) of closely related flaviviral NS4B proteins, according to which 131-169 residues of ZIKV NS4B protein forms the cytosolic region (refer to **figure S7** for MSA of full-length protein). **Figure 5A** shows a part of sequence alignment of NS4B proteins done using Clustal Omega.

**Figure 5:**
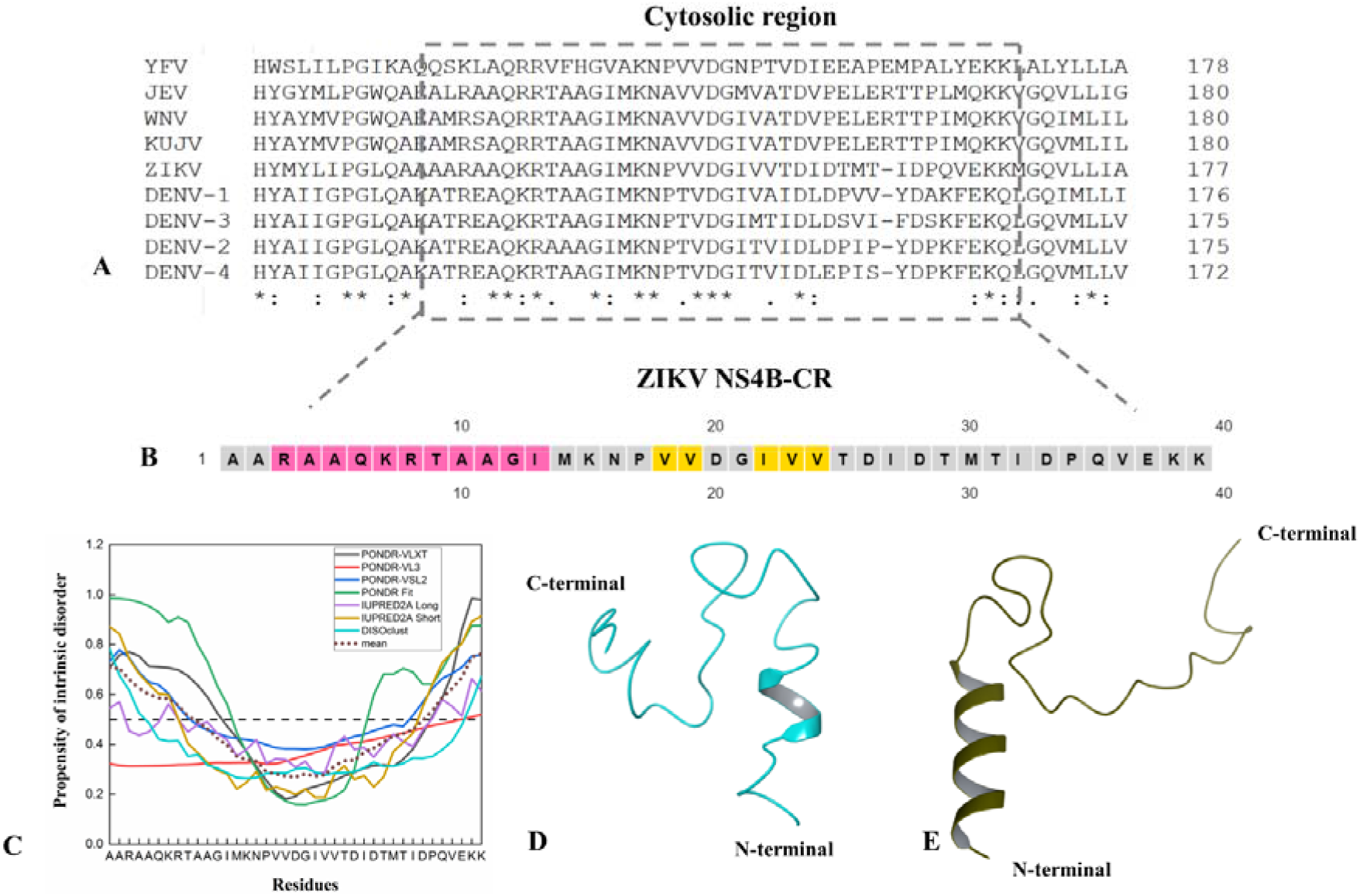
Characterization of the cytosolic region in ZIKV NS4B protein. **(A)** MSA of flaviviral NS4B-FL proteins reveals the cytosolic region forming residues in the ZIKV NS4B protein. (*) represents the conserved residues, and (:) indicates the substitution of amino acid with the residue having strongly similar properties. **(B)** The secondary structure of NS4B cytosolic region (NS4B-CR) predicted using the PSIPRED server shows the presence of residues having helix (pink), β-sheet (yellow), and disordered (grey) propensity. **(C)** Disordered propensity in residues of NS4B-CR is predicted using a combination of PONDR-VLXT, PONDR-VL3, PONDR-VSL2, IUPRED2A long, and short, DISOclust servers and a meta-predictor PONDR-FIT. The mean propensity of intrinsic disorder is depicted with a brown colored dashed line. **(D)** and **(E)** shows the I-TASSER (represented with cyan color) and AlphaFold2 (represented with dark olive color) predicted models of NS4B-CR peptide.

Next, we predicted the secondary structure of NS4B-CR using PSIPRED server which exposed the amino acids having α-helical and -sheet forming propensity (**Figure 5B**). Afterward, we determined the disorder propensity in the cytoplasmic region as it remained disorder as a part of NS4B-FL (**Figure 2**). We employed predictors like the PONDR family, IUPred platform, DISOclust, and a meta predictor PONDR-FIT for this purpose [28–31] (Graph in **figure 5C**). Although the mean susceptibility of intrinsic disorder in middle residues is below the threshold, NS4B-CR has a considerable number of disordered residues. Furthermore, **figures 5D** and **5E** represent the I-TASSER, and AlphaFold2 predicted models of ZIKV NS4B-CR where both modelers revealed the presence of an α-helical region at its N-terminus with an otherwise randomly coiled structure.

#### 3.1. MD simulation of NS4B cytosolic region (NS4B-CR)

We performed 1 μs long simulations of its predicted models (I-TASSER and AlphaFold2) to analyze changes occurring in secondary and tertiary structures (trajectories are shown in **supplementary movies 6** and **7**). It is clear from the captured frames at every 100 ns of the simulation trajectories shown in **figure 6** that NS4B-CR is disordered in isolation. As the trajectory time increases, both modeled structures A and B attained the complete unstructured state. Although for model A, around 12 amino acids at the N-terminal region forming a prominent helix lost their helicity during the simulation in aqueous condition and attained a disordered conformation. The atomic distance-based analysis is illustrated in **figures S8** and **S9**, depicting the increasing RMSD values up to ~10.3 Å and ~6.6 Å for model A and B, respectively. Such high RMSD values with an upward plot trend demonstrate that NS4B-CR is highly dynamic and frequently loses its structure. Additionally, the RMSF values are also higher, suggesting the stability in structure is greatly compromised (**Figures S8** and **S9**).

**Figure 6:**
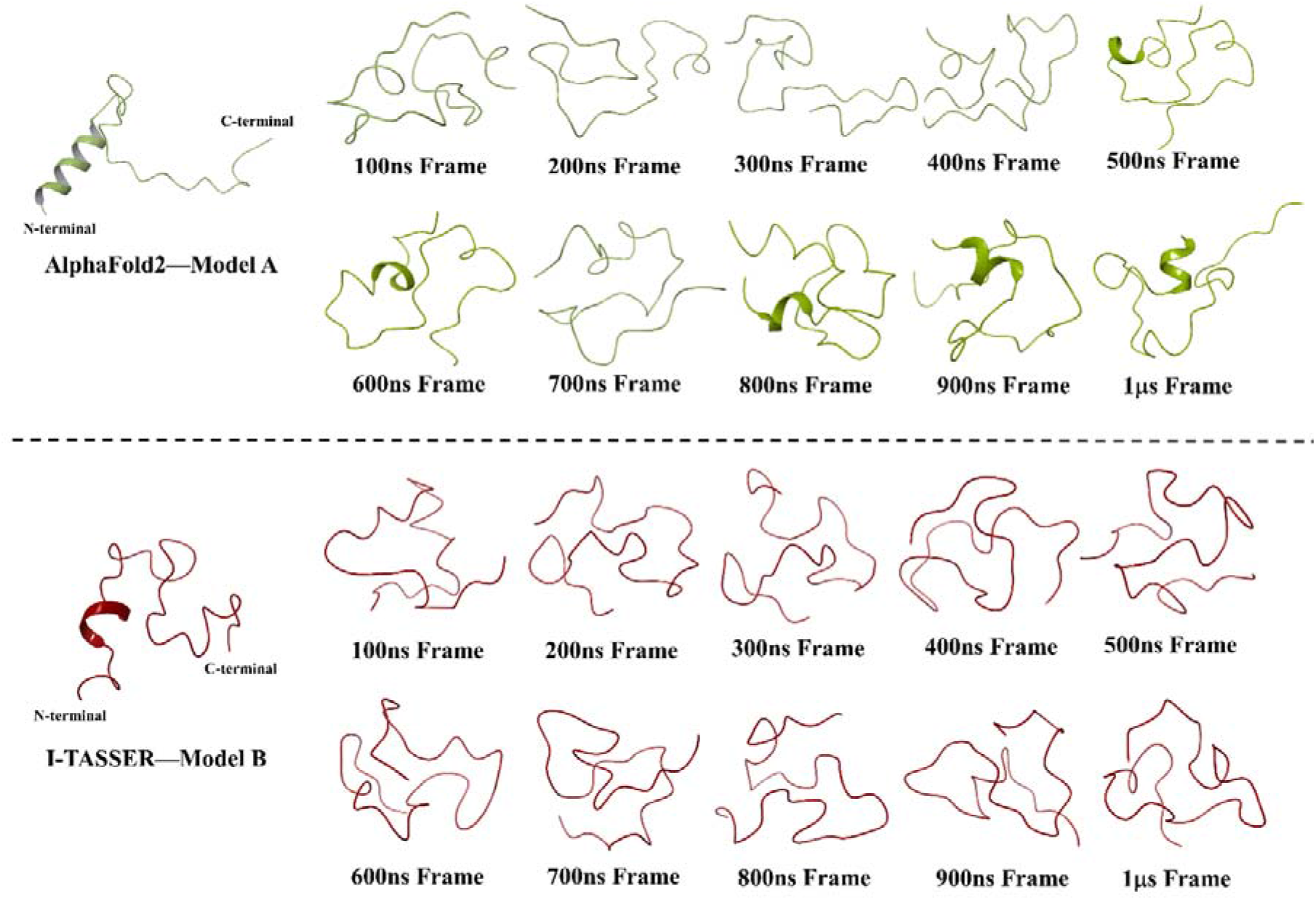
The captured frames of ZIKV NS4B-CR model A (AlphaFold2 model; upper panel; represented with olive color) and model B (I-TASSER model; lower panel; represented with maroon color) at different timeframes during the 1 μs simulations exposed the disordered nature of the cytosolic region.

Moreover, using RMSD based clustering method, all frames of the 1 μs simulation trajectory of model A were analyzed and found to correlate with the above analysis. The top 10 structures, which represent a cluster of similar structures, are shown in **figure S10**. All structure representatives of model A have RMSD values ranging between ~6.3-7.3 Å.

#### 3.2. REMD simulation analysis

After intrinsic disorder prediction and simulation of ZIKV NS4B-CR in isolation which nevertheless remains a useful strategy for characterizing an IDP or IDPR, we additionally performed a REMD simulation using model B structure (I-TASSER predicted model). The snapshots at 100 ns and 200 ns in inner and outer circles shown in **figure 7**, respectively, provide a clear idea of the intrinsic property of disorderedness of NS4B-CR. At lower temperatures, there is no significant change observed in the structure. However, at temperatures like 330K, 350K, 370K, 400K, there is a gain of one or two β-sheet regions, present in either in one or both represented frames of replicas, which were nonetheless reversible. This may signify that the exchange between parallel replicas could not be feasible, and hence the peptide is not susceptible to gain structure easily. Likewise, the secondary structure timelines (shown in **figure S13**) also emphasize the fact that NS4B-CR is not able to gain any stable structure in isolation and at any temperature (except a few frames). The overall changes in secondary structure elements (SSE) are shown in **supplementary table 1**, where less than 5% of the total secondary structure is formed throughout the simulation period. Furthermore, the statistical analysis based on mean atomic distances (RMSD and RMSF plots in **figures S11** and **S12**) evidently display the flexible nature of NS4B-CR.

**Figure 7:**
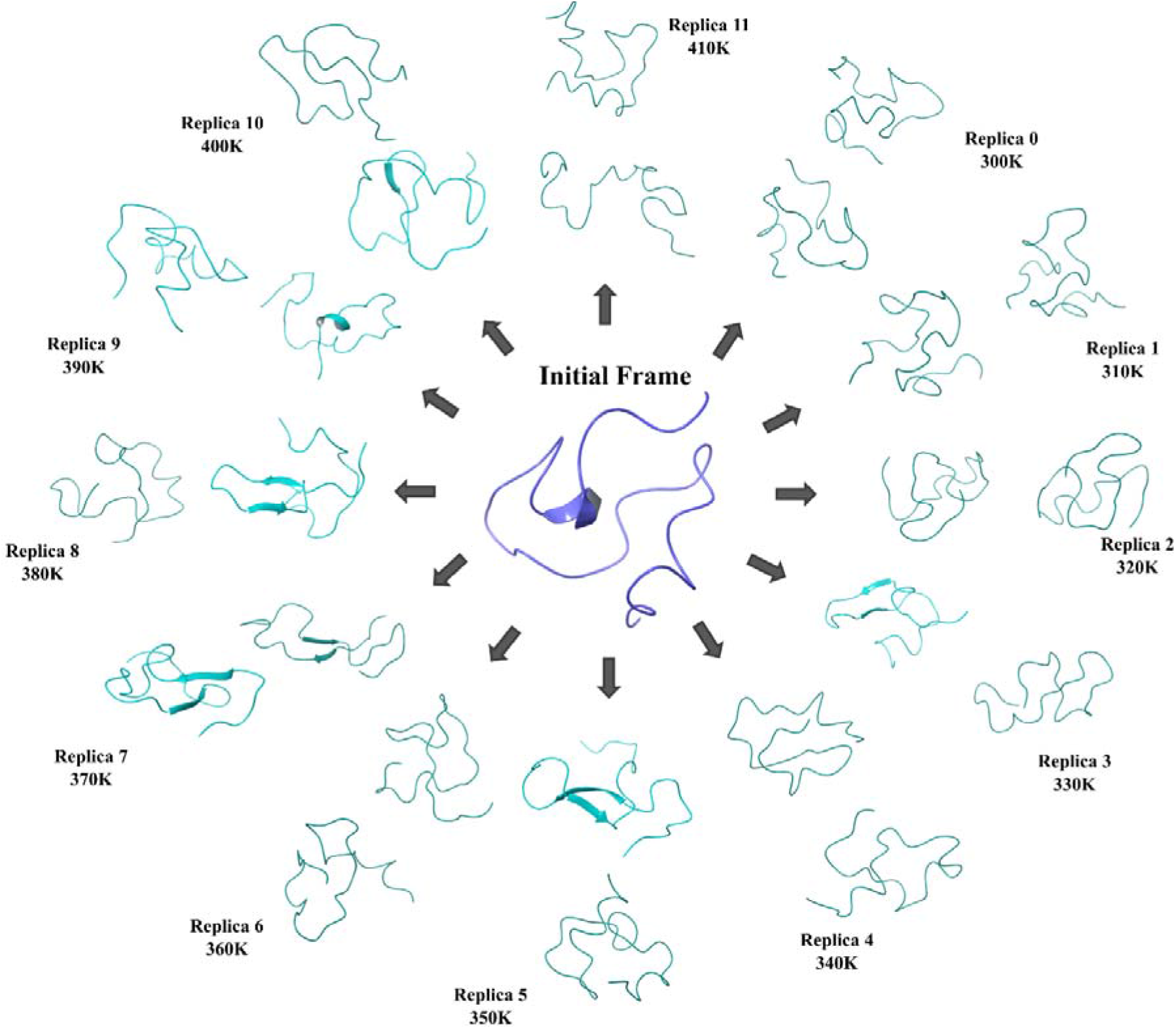
Depiction of snapshots from all replicas (0-11) at every 100 ns (the inner circle) and 200 ns (outer circle) period during the REMD simulation of NS4B-CR.

### 4. Experimental validation of unstructured nature of NS4B-CR and propensity of “gain-of-structure”

#### 4.1. Disordered nature of NS4B-CR

We experimentally validated NS4B-CR peptide to remain in disordered conformation in buffered conditions. Far-UV CD spectrum where negative ellipticity near 198 nm is indicative of its flexible nature is shown in **figure 8A**. Besides, no other characteristic peak showing the presence of α-helix or -sheet is found in the CD spectrum. Further, MoRFs in the cytosolic region are confirmed by observing its “gain-of-structure” potential in the presence of solvents providing distinct environmental conditions. MoRFs are segments within the disordered protein region which undergo disorder-to-order transition. They serve as initial contact points for interaction events that concomitantly gain structure on induced binding by any physiological partner. Firstly, the behavior of NS4B-CR peptide under the influence of TFE, a secondary structure inducer, was examined [32]. As anticipated, on increasing the concentration of TFE, it gradually forms a helical structure. Pronounced negative ellipticity values at 206 nm and 222 nm in CD spectra in 70% TFE indicate the formation of α-helical conformation in the previously disordered peptide (**Figure 8C**). **Figure 8D** represents the relationship of ellipticity values at 198 nm and 222 nm with TFE concentration. Similarly, in the presence of TMAO, an osmolyte [33], NS4B-CR peptide gains a partial helical conformation (**Figure 8B**).

**Figure 8:**
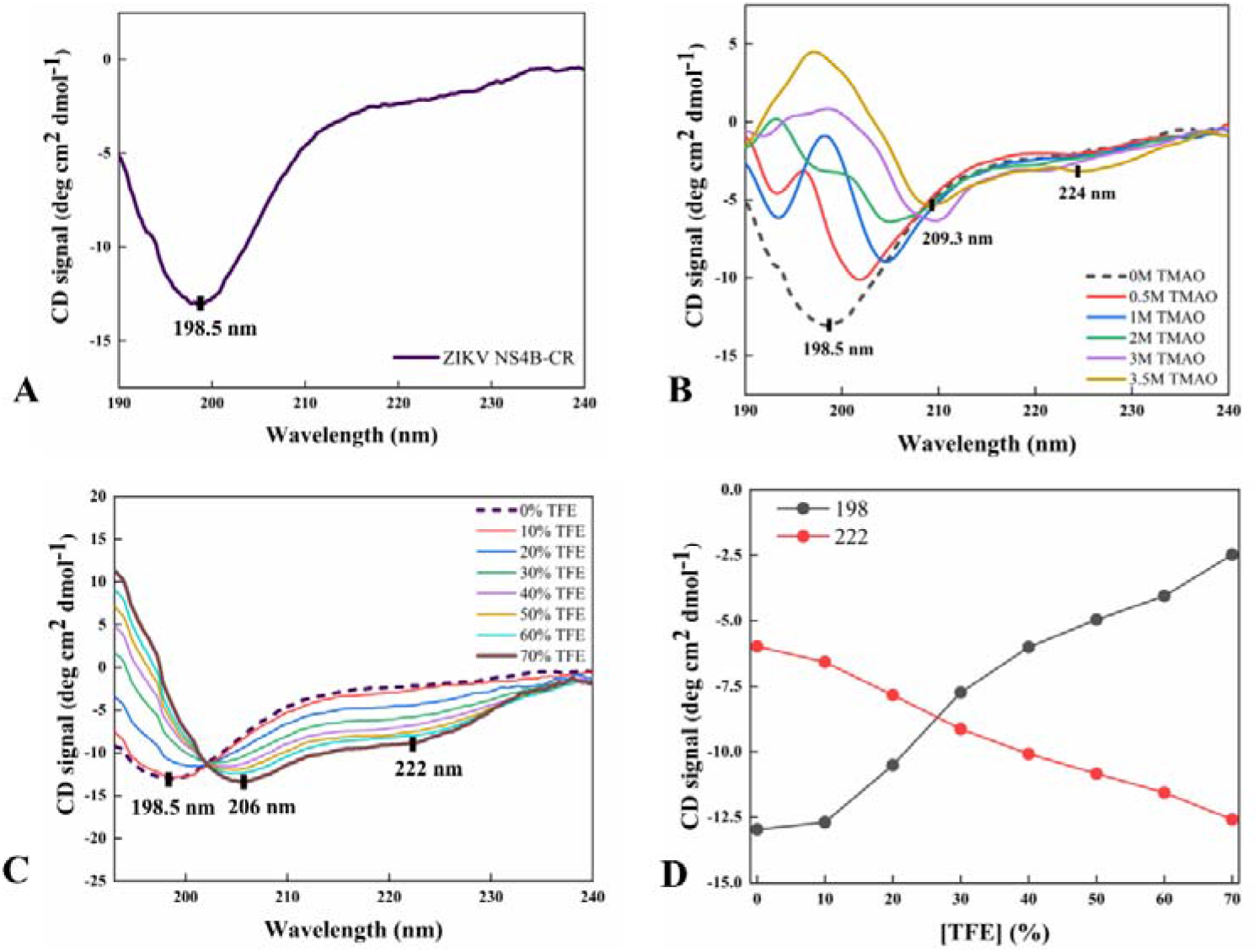
Experimental validation of unstructured nature of ZIKV NS4B-CR (131-169 residues). **(A)** The prominent negative peak of NS4B-CR far-UV CD spectra at 198.5 nm reveals the disordered conformation of this region experimentally. Spectral changes in the presence of **(B)** TMAO, an osmolyte, and **(C)** TFE, a known secondary structure inducer. **(D)** Effect of increasing concentration of TFE on NS4B-CR far-UV CD spectra at 198 nm and 222 nm.

#### 4.2. Membrane-mediated effects on NS4B-CR

To analyze the ability of NS4B-CR peptide to interact with cellular lipid molecules, we studied its behavior in the presence of lipids and artificial membrane forming SDS micelles [34]. As shown in **figures 9A** and **9B**, the maximum negative ellipticity of CD spectra of NS4B-CR peptide under the influence of SDS micelles shifts from wavelength 198.5 nm to 201.6 nm. This indicates the formation of a pre-molten globule structure in SDS. The same has been confirmed by analyzing the CD spectra using an online server known as CAPITO (**Figure S14**) [35]. However, in the presence of liposomes: DOPC – neutrally charged; DOPS – negatively charged; and DOTAP – positively charged, NS4B-CR remains disordered and does not show any charge-based interaction with liposomes (**Figures 9C-H**). These results suggest that either NS4B-CR does not interact with lipid molecules or remains disordered after the interaction.

**Figure 9:**
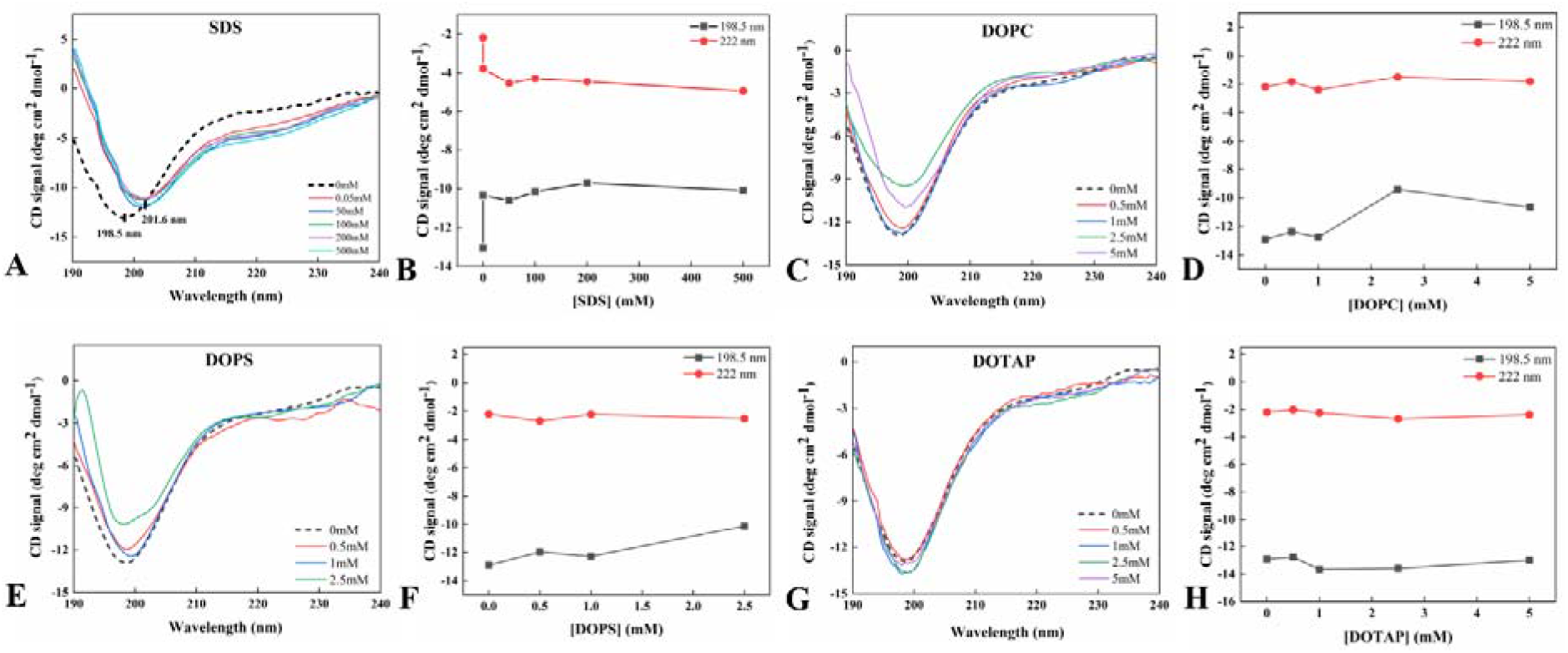
Membrane-mediated effects on NS4B-CR. (**A**) Changes in the secondary structure of NS4B-CR due to SDS micelles. (**B**), (**C**) and (**D**) represents the conservation of disordered nature of NS4B-CR peptide in the presence of DOPC (neutrally charged), DOPS (negatively charged), and DOTAP (positively charged) liposomes, respectively.

#### 4.3. Behavior of NS4B-CR under macromolecular crowding

Macromolecular crowding explains the substantial nonspecific influence of volume excluded by one molecule to another. It mimics the cellular environment where the total concentration of macromolecules can reach up to 80-400 mg/ml [36,37]. Thus, we have investigated the influence of well-known crowding agents like Ficoll-70 and Poly Ethylene Glycol 8000 (PEG8000) on the NS4B-CR structure. As shown in **figures 10A**, **10B**, **10D**, and **10E**, no considerable effect of crowding is observed on its structure as no significant change in structure is detected in the representative far-UV CD negative peak at 198 nm.

**Figure 10:**
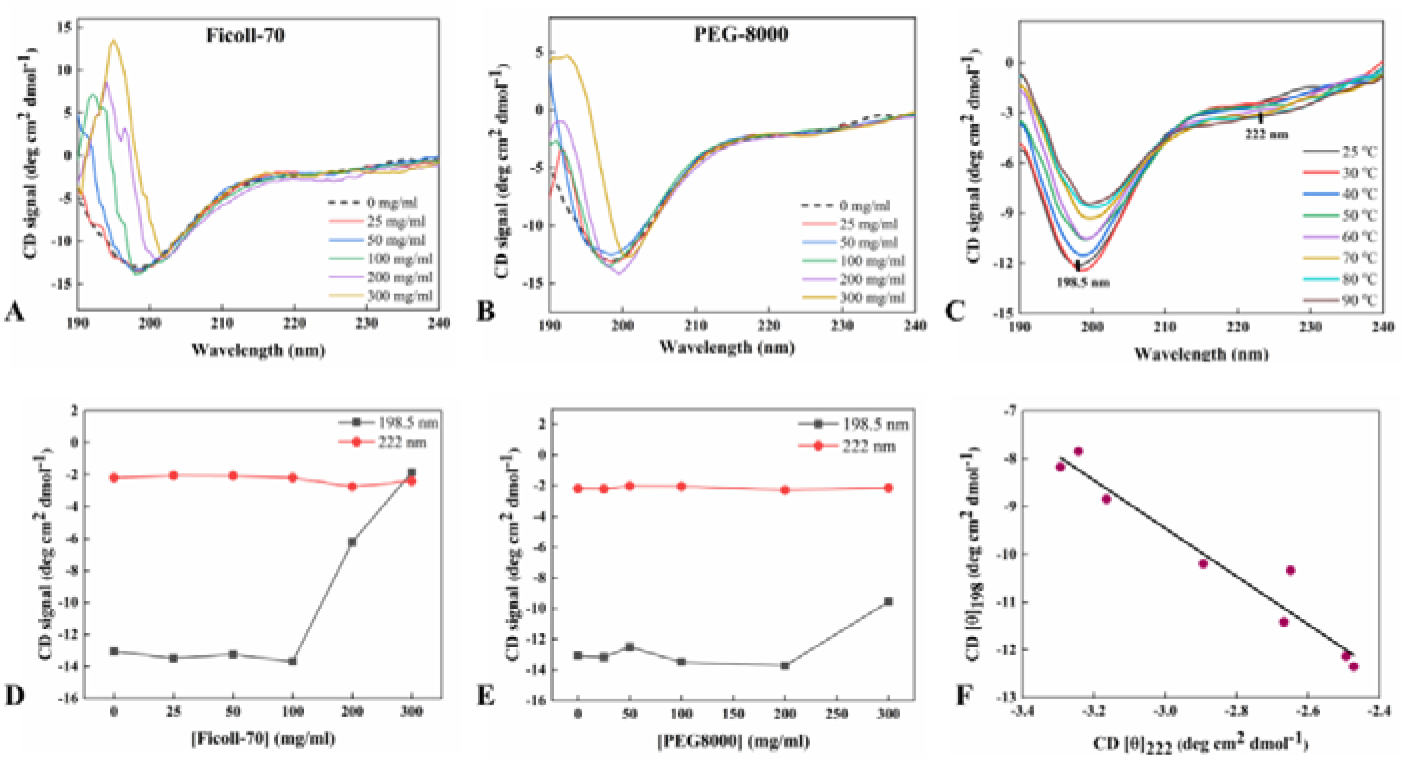
Macromolecular crowding and temperature-mediated effects on disordered NS4B-CR region. **(A)** and **(D)** represents the changes occurring in far-UV CD spectra of NS4B-CR in the presence of Ficoll-70, **(B)** and **(E)** shows similar changes in CD spectra in the presence of PEG8000. The graph in **(C)** depicts the decrease in negative ellipticity values at 198 nm of CD spectra on an increment of temperature. **(F)** The phase diagram was plotted using ellipticity values at 222 nm on the x-axis and 198 nm on the y-axis. The straight line represents the two-state folding mechanism.

#### 4.4. Effect of variable temperature on NS4B-CR

Extended IDPs, in general, acquire a “turned out” response to heat where partial folding of certain proteins has been observed [19,38]. The limited folding in IDPs and IDRs is attributed to an increase in intramolecular hydrophobic forces. Similarly, increasing temperature induces partial folding of NS4B-CR peptide. However, it should be noted that, as such, there are no significant conformational changes. The slight changes in the CD spectra suggest a typical temperature-induced behavior of IDPs known as contraction. These structural changes shown in **figures 10C** and **10F** represents the strengthening hydrophobic interactions leading to incomplete folding. Furthermore, spectral changes are analyzed using the phase diagram where [0]_198_ are plotted against [0]_222_ (**figure 10F**). The straight line in a phase plot reflects an all-or-none transition in the folding pathway while a break in linearity represents the presence of an intermediate state [39]. Thus, in the NS4B-CR peptide, the partial compactness towards a more ordered structure occurs using a two-state folding mechanism.

## Discussion

Despite the advancements in techniques and approaches, the biophysical studies establishing the topology of membrane proteins are frequently less performed due to their hydrophobic character. Biochemical studies on flaviviral NS4B proteins have not become a central focus like the flaviviral NS3 helicase and NS5 RNA-dependent RNA polymerase proteins. Moreover, it has become important to investigate the dynamicity of its transmembrane and non-membranous regions in isolation to understand structural transitions. So far, only DENV2 and WNV NS4B proteins have been studied for their topology [7–9]. Herein, we studied the structural dynamics of full-length NS4B protein of ZIKV and its three significant non-membranous regions in isolation.

N-terminal region of flaviviral NS4B protein is involved in host IFN pathway antagonism. This function of NS4B protein is conserved in closely-related flaviviruses - YFV, WNV, and DENV [2]. In WNV, NS4B-P38G mutant produces lower viremia, no lethality, and higher IL-1β and IFNs responses in mice as compared to wildtype strain [40]. In human cell lines THP-1 cells and THP-1 macrophages, the same mutant boosted the innate cytokine responses [41]. According to our results, ZIKV NS4B N-terminal region has a tendency to form disordered structure and demonstrates dynamicity as a part of full-length protein. In isolation, it completely changes its conformation to unstructured after 1μs of simulation.

Further, C-terminal region which contains a membrane-spanning region according to literature as well as predictions. In microsecond simulations, it loses structure in absence of POPC membrane, but in presence of the membrane, the model retained its original secondary structural conformation in isolation. Moreover, as a part of full-length NS4B, the C-terminus residues show substantial fluctuations leading to high RMSF values as compared to other membrane interacting residues. This shows that C-terminal region may not have significant hydrophobicity to span the membrane completely, and thus on residing in ER membrane, it remains intact while after cleavage from NS5 protein, it attains disordered conformation in ER lumen.

DENV and WNV NS4B proteins are membranotropic in nature containing a loop of ~ 35 amino acids between transmembrane domains 3 and 4 [8]. This protein segment residing in the cytoplasm, generally called a cytosolic loop, is assumed to be essential for the whole protein. Characterizing the structural state of this region in ZIKV NS4B protein is of great importance. According to the conventional model of flaviviral replication site, this region of NS4B protein lies in the cytoplasmic part of virus-induced membranous regions [42]. DENV NS4B protein via its cytosolic loop interacts with NS3 protein in virus-induced sites and acts as a cofactor for its helicase activity where it binds to subdomains 2 and 3 of the helicase domain of NS3 protein with an affinity of 340 nM [11]. Secondly, the cytosolic loop appears to initiate the dimerization of two NS4B proteins in DENV, which gives us another reason for investigating the structural dynamics of this region in ZIKV [9]. Therefore, through multiple sequence alignment of different flaviviral NS4B proteins, we first identified the residues forming the similar cytosolic region (CR) in ZIKV NS4B protein. Then, we performed CD spectroscopy-based experiments with the synthesized peptide (NS4B-CR) to check its secondary structure conformations in isolation and by providing different environments in-vitro. The signature peak in CD spectra between 195 nm - 200 nm demonstrates it to be a disordered region in isolation and no significant shift in wavelength in membrane-mimetic and crowding conditions also suggest the same. In addition, the results from microsecond long MD simulations of both I-TASSER and AlphaFold2 predicted models of NS4B-CR established it to be intrinsically unstructured. Based on our overall observations, we have proposed a putative model of NS4B protein (**Figure 11**) that illustrates the possible architecture of transmembrane and non-transmembrane regions of NS4B in Zika virus. Specifically, for C-terminal region, we have identified that it forms ordered and disordered conformations in presence and absence of lipid bilayer, respectively. This reflects that it may have the tendency to remain functional in membrane as well as in lumen through its structural transitional ability.

**Figure 11:**
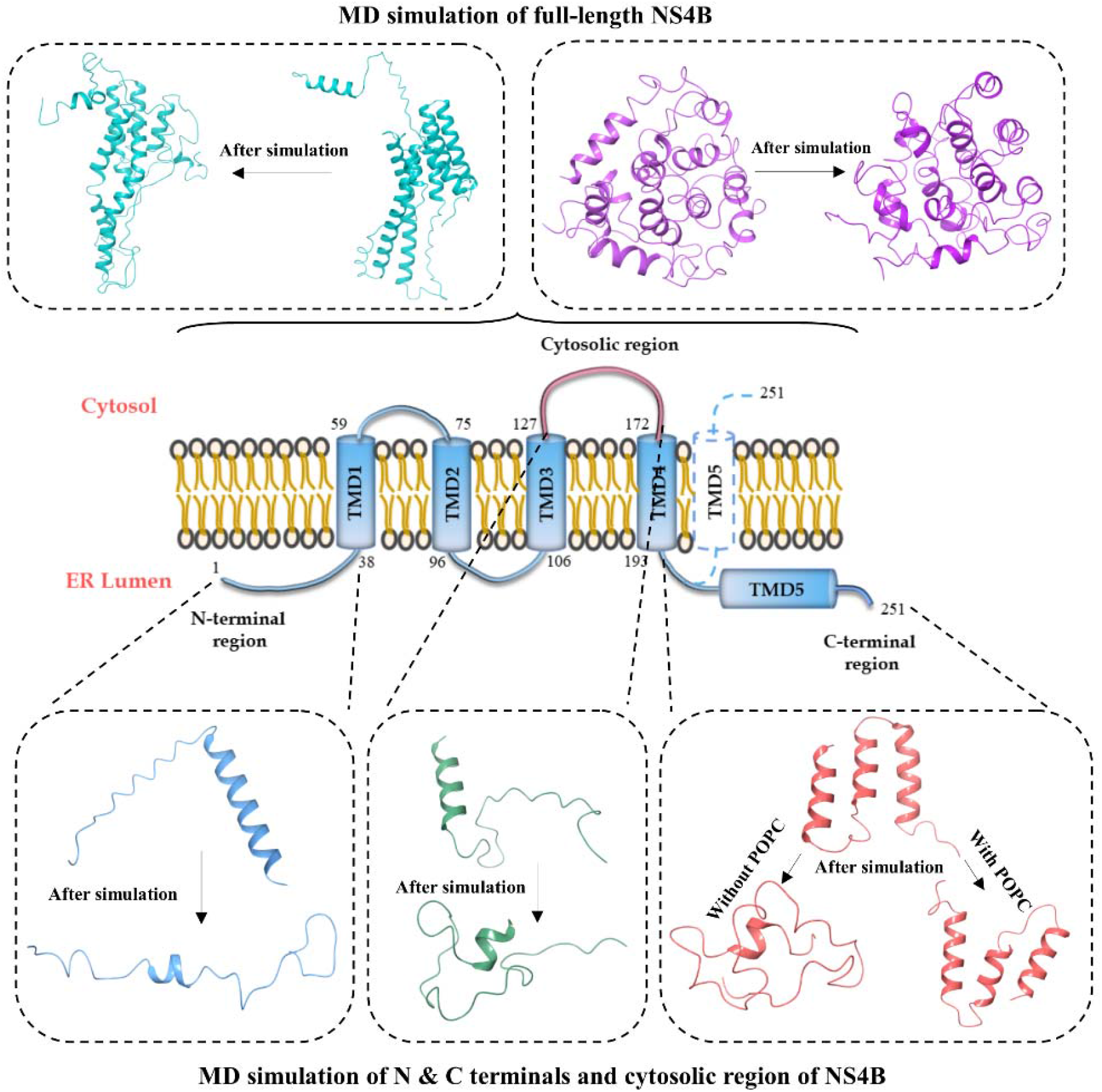
Proposed model of ZIKV NS4B protein and representation of transmembrane and non-membranous regions using 3D structure models before and after MD simulations.

A large body of literature suggests that terminal regions of viral proteins have considerable amount of disorder in them. This renders them to undergo conformational rearrangements on interaction with their physiological partner. For instance, ZIKV capsid and NS4A proteins, have disordered N-terminal regions [17,43]. The N-terminal residues 1-48 of ZIKV NS4A further gains secondary structure on interacting with liposomes [43]. In SARS-CoV-2, C-terminal region of spike protein is analyzed to remain in disordered conformation [27]. Nucleocapsid protein of coronaviruses also have disordered N- and C-terminals [44,45]. N-terminal half of core protein of HCV (C82) is mostly unstructured and is a well-documented IDP region [46]. Similar examples of unstructured terminal regions of proteins are present in different viruses which clearly reflect the functional relevance of such regions [47,48].

Likewise, the transmembrane proteins have preference for structural disorder at their terminals and loops. These disorder-rich terminal and loop regions possess the ability to mediate interactions with external partners and can even send regulatory signals within the protein [49]. However, the distribution of IDPRs is five times more in terminals than in loop segments [49]. The disordered regions which are substantially more abundant in residues like valine, isoleucine, and alanine for their inherent sensitivity to fold on interaction with surrounding molecules allows them to gain structure. The flexible nature of unstructured terminals and loops renders them to typically undergo rapid unimolecular steps and permit the creation of stable complexes with a number of targets through the fly-casting mechanism [50]. In such scenario, IDPRs have a large capture radius available for the target site interaction at the cost of entropy where it is traded off with the energy gained upon binding to the target protein alongside gaining a secondary structure [50,51]. Earlier, certain MoRF regions in ZIKV NS4B protein were identified from residues 1-10, 33-37, and 241-250 [52]. Correspondingly, the highly dynamic regions of NS4B protein may assist them in interaction with other viral and host protein.

## Conclusion

Previously, studies have experimented with the full-length flaviviral NS4B proteins, however, none of them commented about the secondary structure which it can adapt inside the cells or in-vitro. Therefore, to extend the understanding in this direction, we have investigated the secondary structural characteristics of three critical non-membranous regions of ZIKV NS4B protein. We have used computational and experimental approaches to show that the cytosolic region of ZIKV NS4B protein is highly dynamic and thus is an intrinsically disordered protein region. It also possesses gain of structure property under specific environment like TFE and TMAO but do not form helical or beta-strands in lipids or macromolecular crowders. Simulations results illustrate that the N-terminal region lose its compact secondary conformations in isolation and also becomes unstructured as a part of NS4B-FL protein. Furthermore, C-terminal region also has the tendency to be dynamic which may assist its interaction with membrane lipids whenever required.

## Authors contribution

RG: Conception and design, review of the manuscript, and study supervision. TB and PK: Performed experiments, writing, and review of the manuscript.

## Acknowledgments

Authors would like to thank IIT Mandi for providing facilities. RG is thankful to SERB, Government of India (Grant no. CRG/2019/56003), MHRD-SPARC (SPARC/2018– 2019/P37/SL), IYBA award from DBT, Government of India (BT/11/ IYBA/2018/06), and ICMR Government of India (Grant no. 58/6/2020/PHA/BMS). TB is grateful to the DST, Government of India, for her INSPIRE fellowship for funding.

## Conflict of interest

All authors declare that there is no competing financial interest.

## Notes

### Competing Interest Statement

The authors have declared no competing interest.

