## Supplementary Table 1, Supplementary Figures for "Investigating the conformational dynamics of Zika virus NS4B protein"

**Supplementary Information**

**Supplementary Figures**

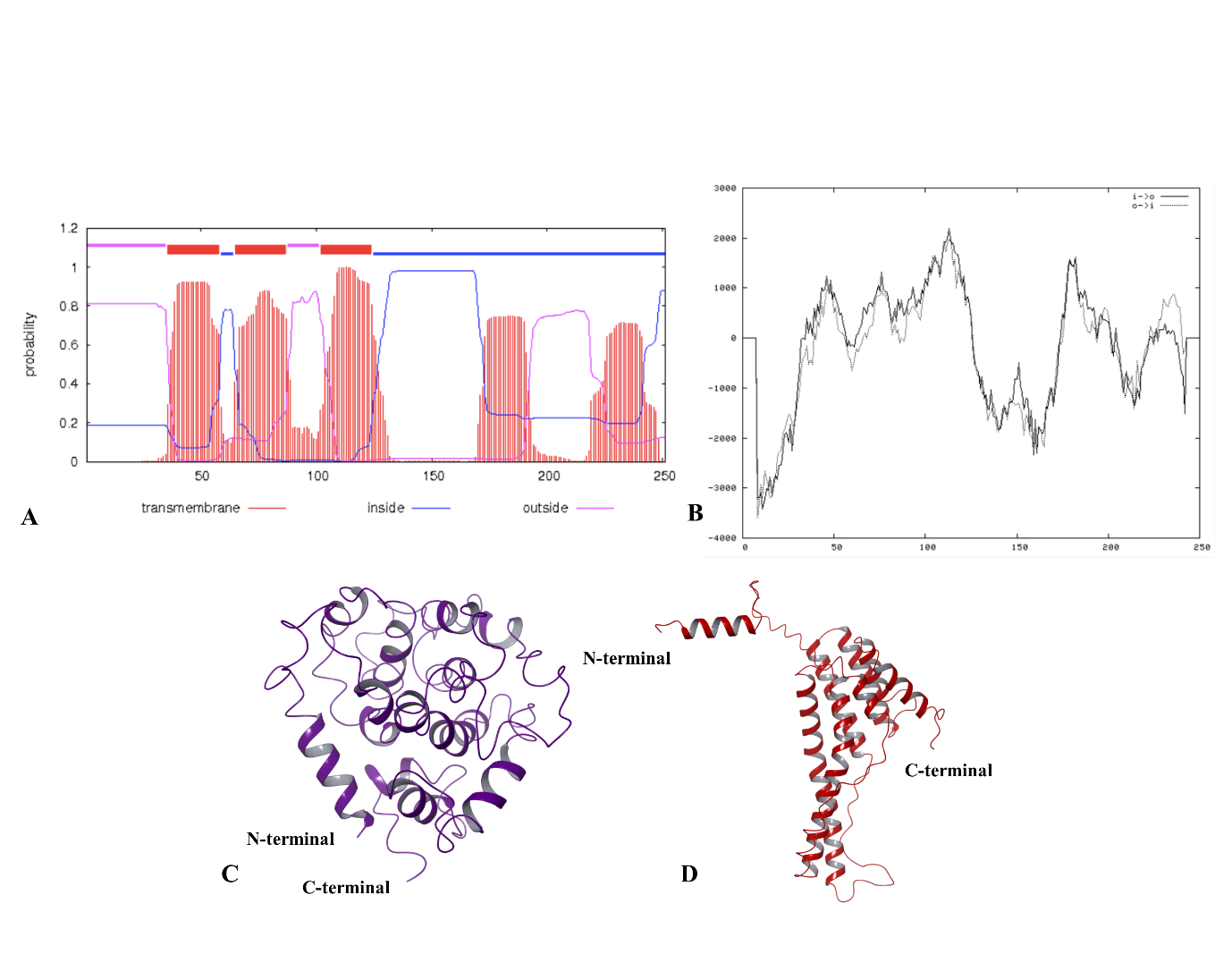

**Figure S1:** The transmembrane regions in ZIKV NS4B are predicted using TMHMM and TMPred server. Accordingly, TMHMM shows three full transmembrane regions while TMPred results shows the presence of five membrane-spanning regions.

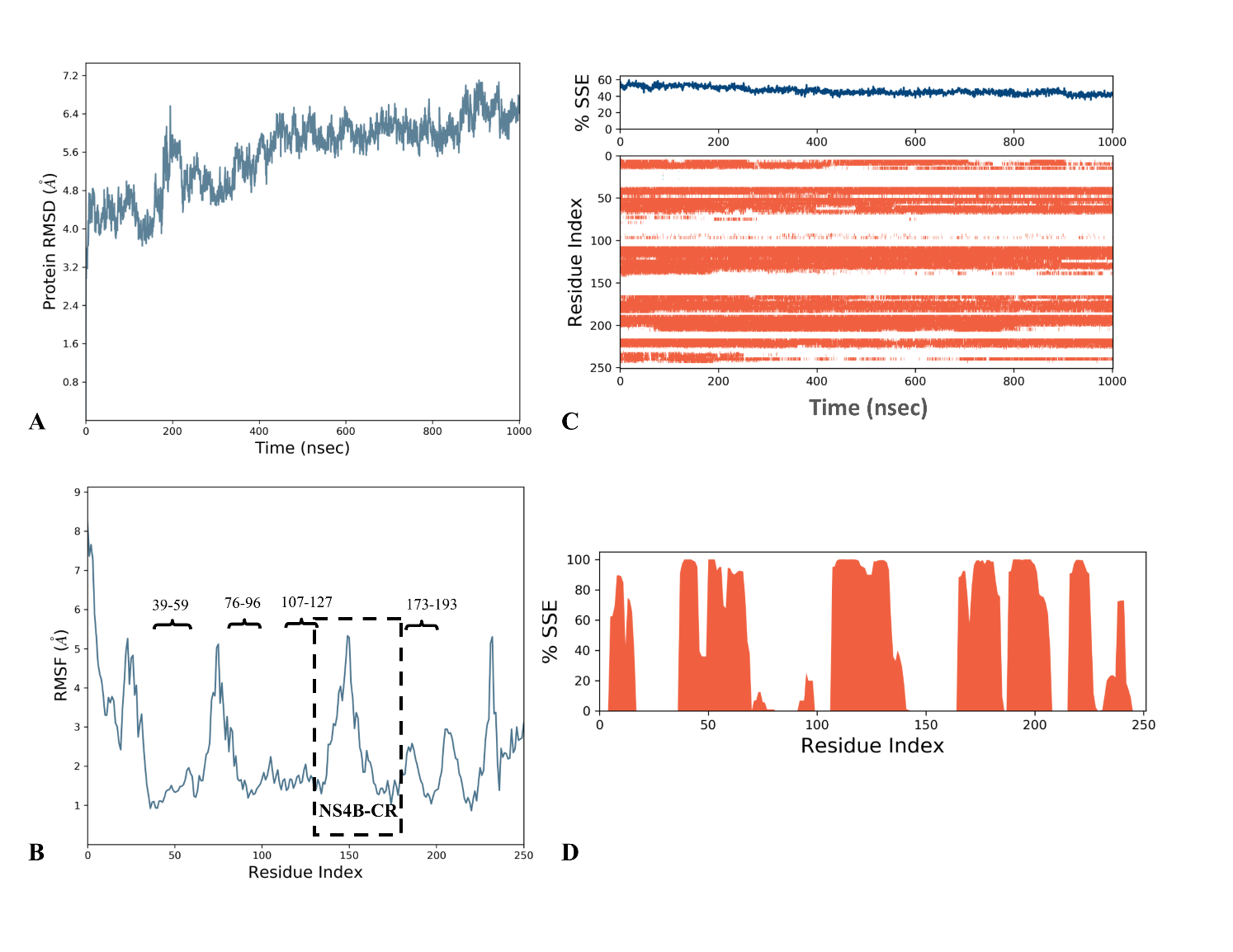

**Figure S2:** **One microsecond simulation of NS4B-FL model A predicted using AlphaFold2**: **(A)** RMSD, **(B)** RMSF, and **(C** and **D)** secondary structure analysis through timeline.

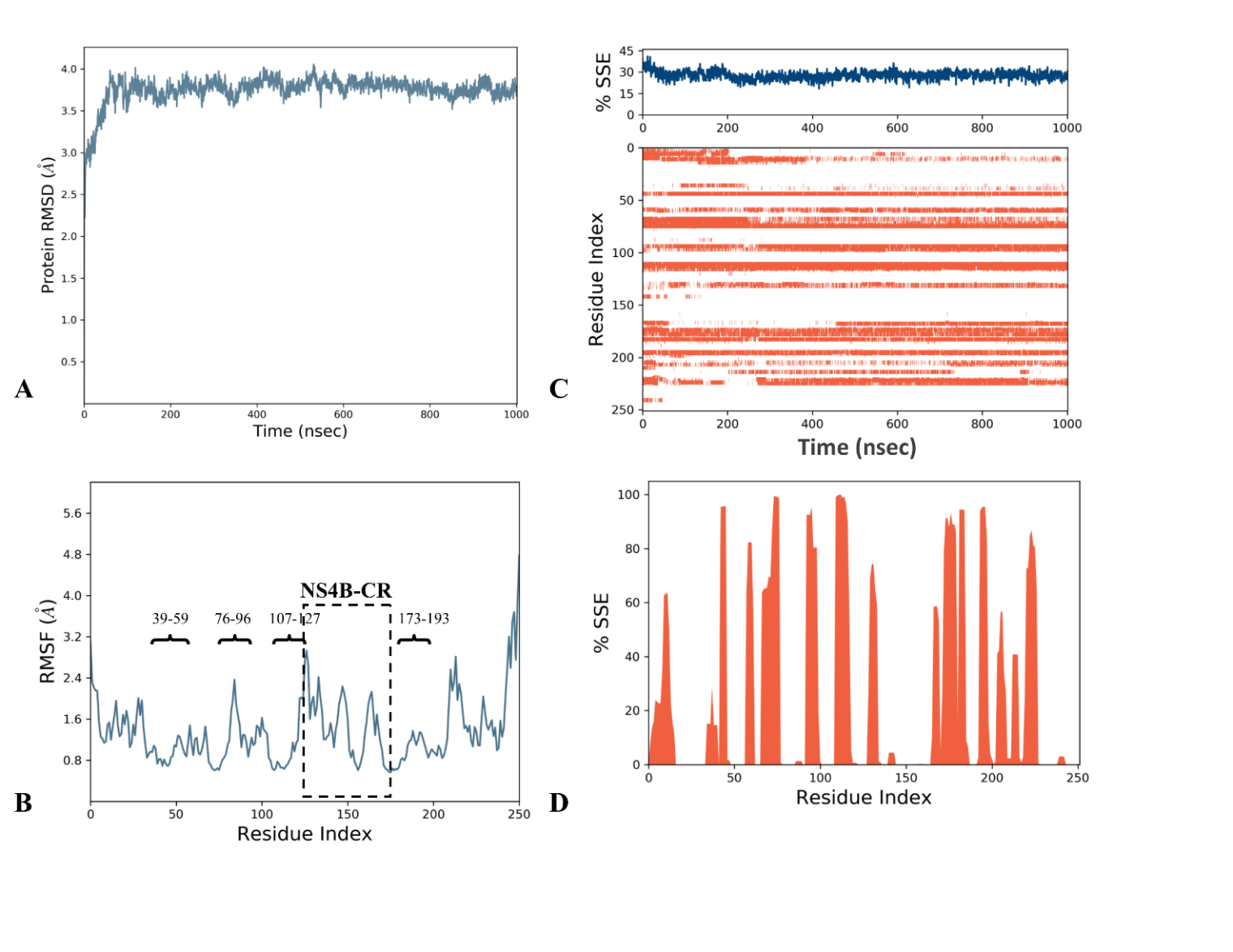

**Figure S3:** **One microsecond simulation of NS4B-FL model B predicted using I-TASSER: (A)** RMSD, **(B)** RMSF, and **(C** and **D)** secondary structure analysis through timeline.

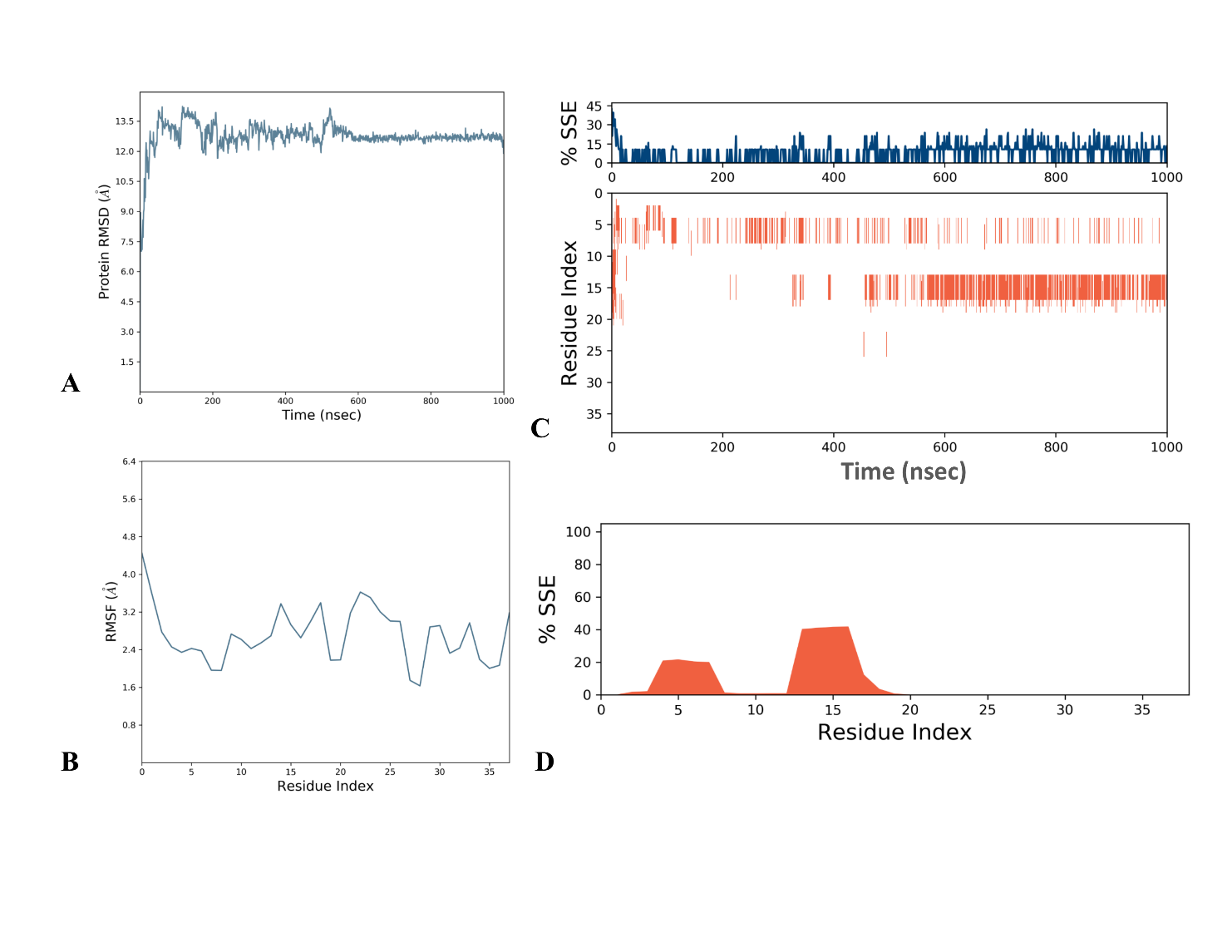

**Figure S4:** **One microsecond simulation of NS4B-NT model predicted using AlphaFold2:** **(A)** RMSD, **(B)** RMSF, and **(C** and **D)** secondary structure analysis through timeline.

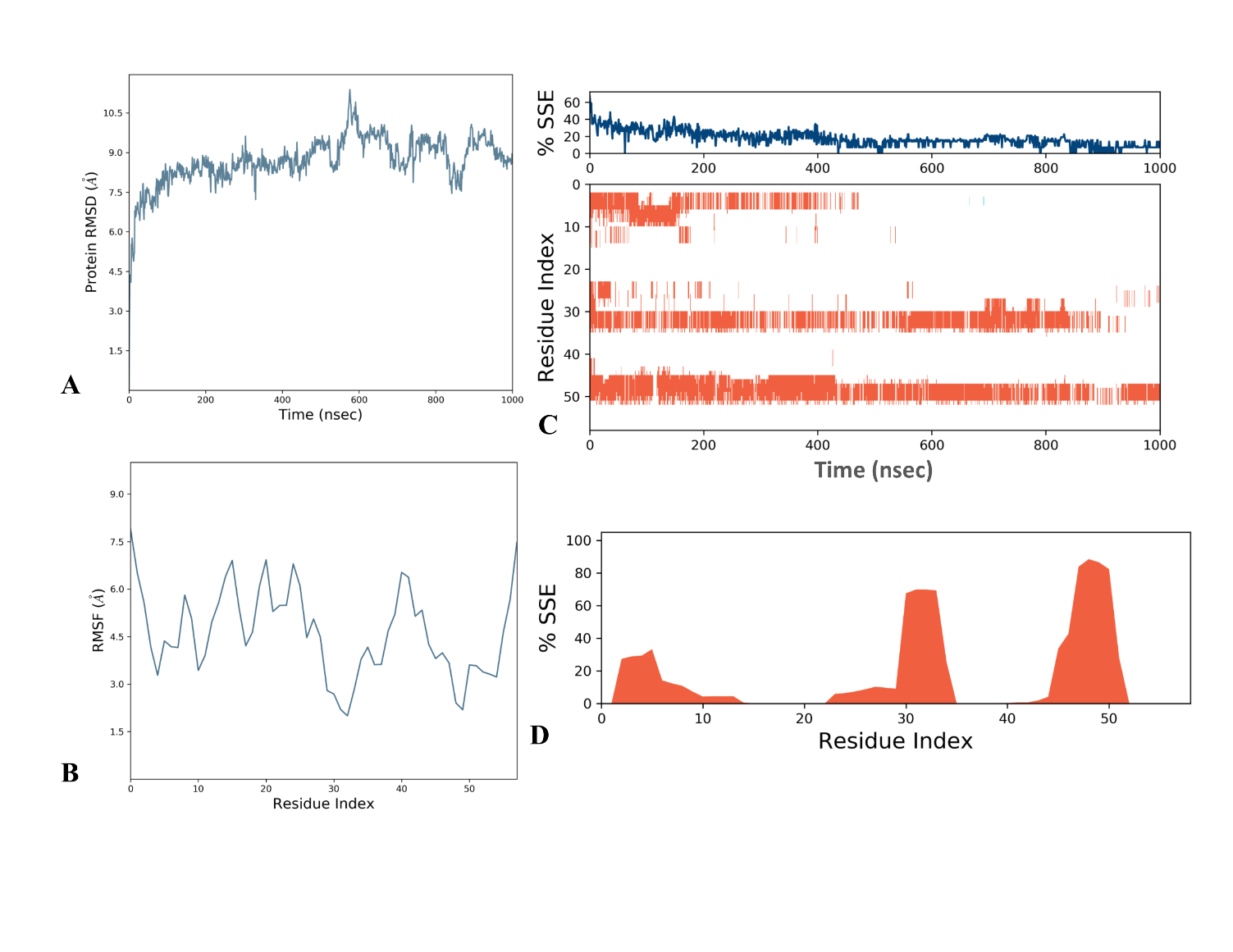

**Figure S5: One microsecond simulation of NS4B-CT model predicted using AlphaFold2 in the absence of POPC membrane:** **(A)** RMSD, **(B)** RMSF, and **(C** and **D)** secondary structure analysis through timeline.

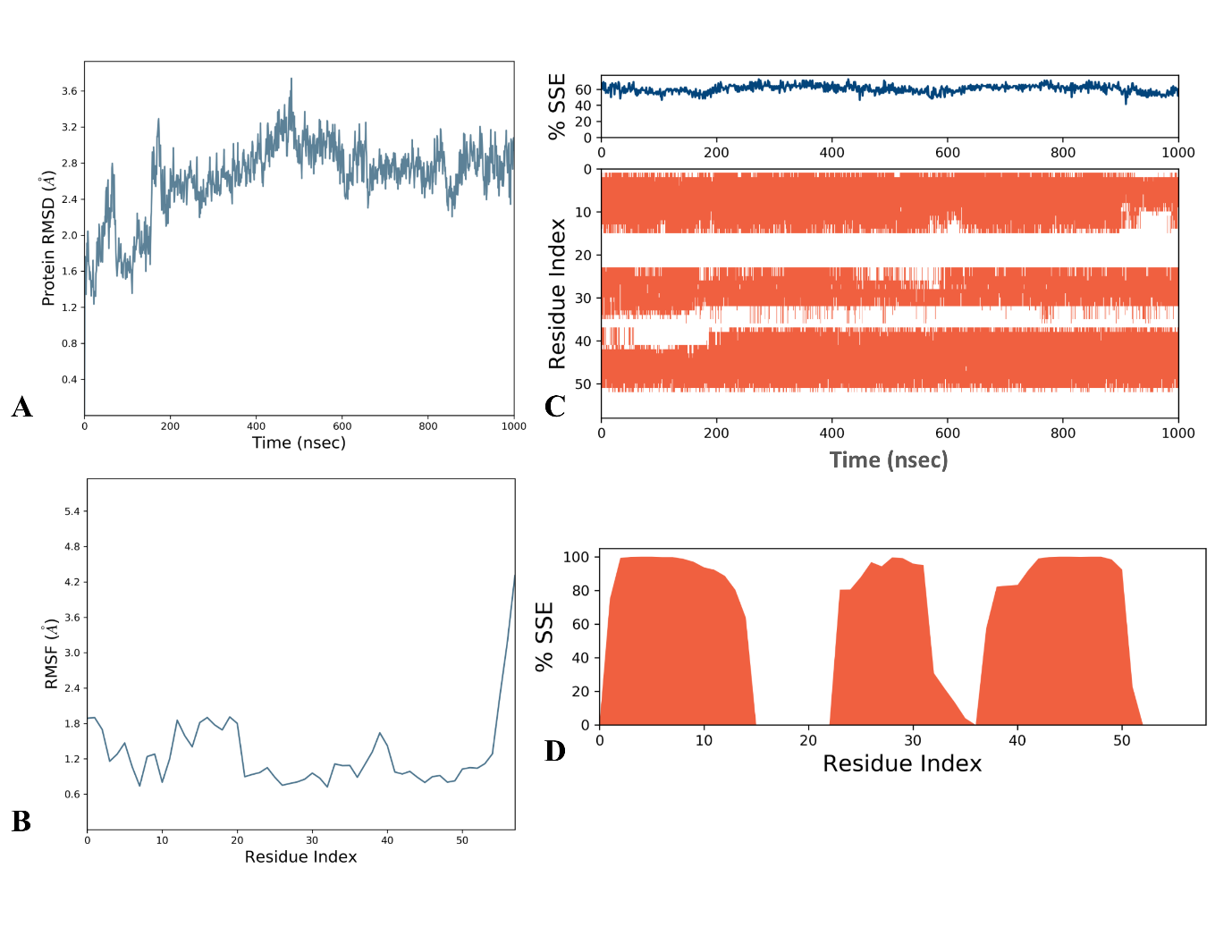

**Figure S6:** **One microsecond simulation of NS4B-CT model predicted using AlphaFold2 in the presence of POPC membrane:** **(A)** RMSD, **(B)** RMSF, and **(C** and **D)** secondary structure analysis through timeline.

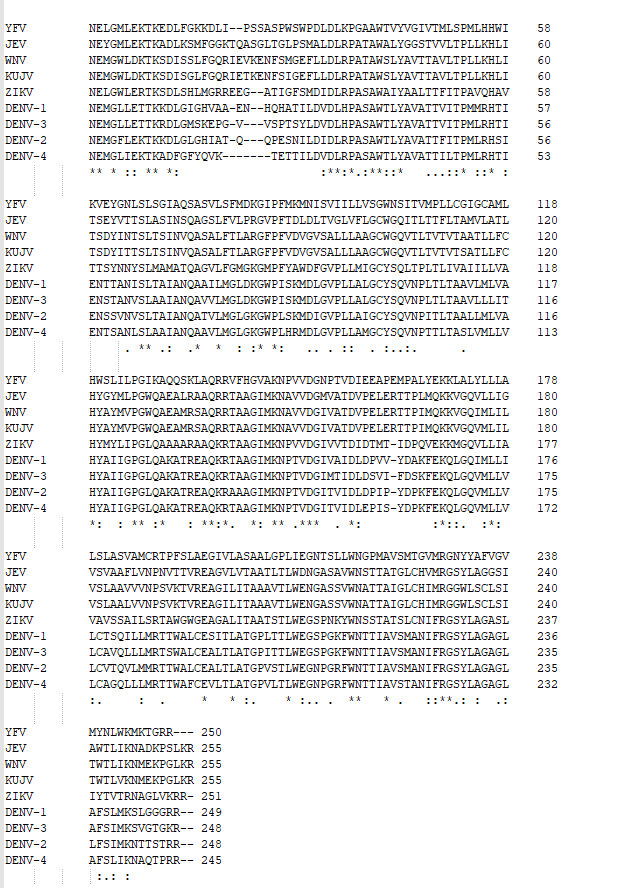

**Figure S7:** Multiple sequence alignment of full-length flaviviral NS4B proteins.

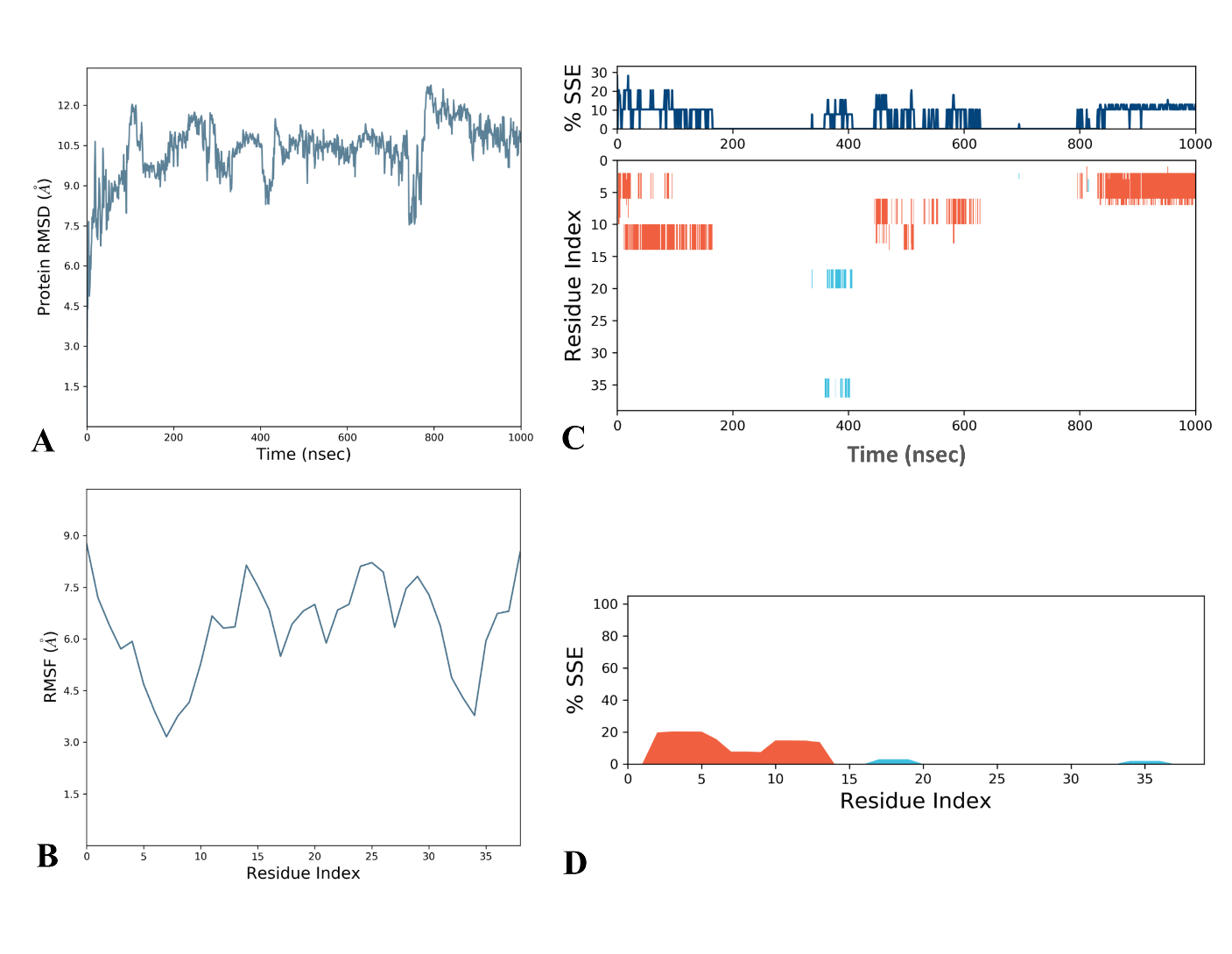

**Figure S8:** **One microsecond simulation of NS4B-CR model A predicted using AlphaFold2**: **(A)** RMSD, **(B)** RMSF, and **(C** and **D)** secondary structure analysis through timeline.

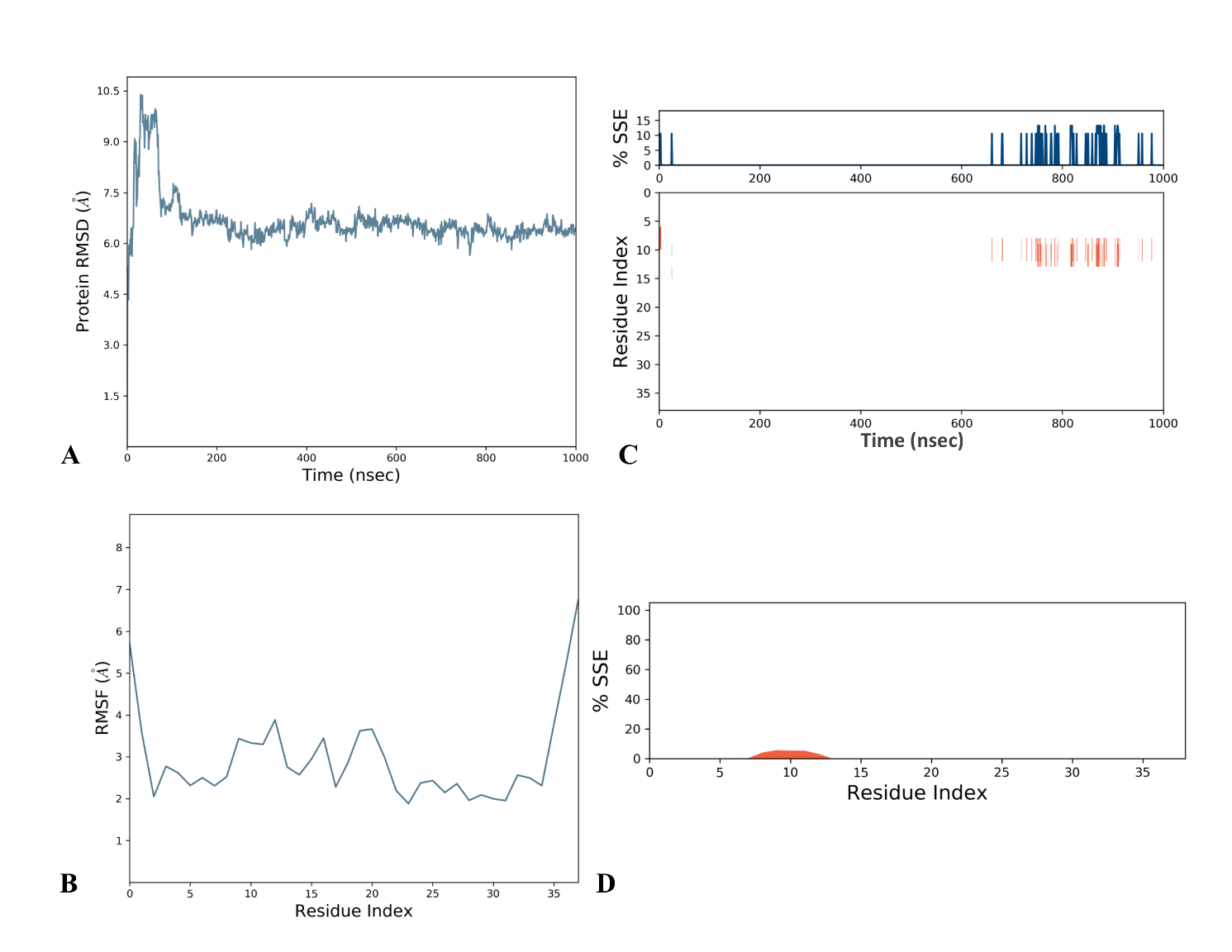

**Figure S9:** **One microsecond simulation of NS4B-CR model B predicted using I-TASSER: (A)** RMSD, **(B)** RMSF, and **(C** and **D)** secondary structure analysis through timeline.

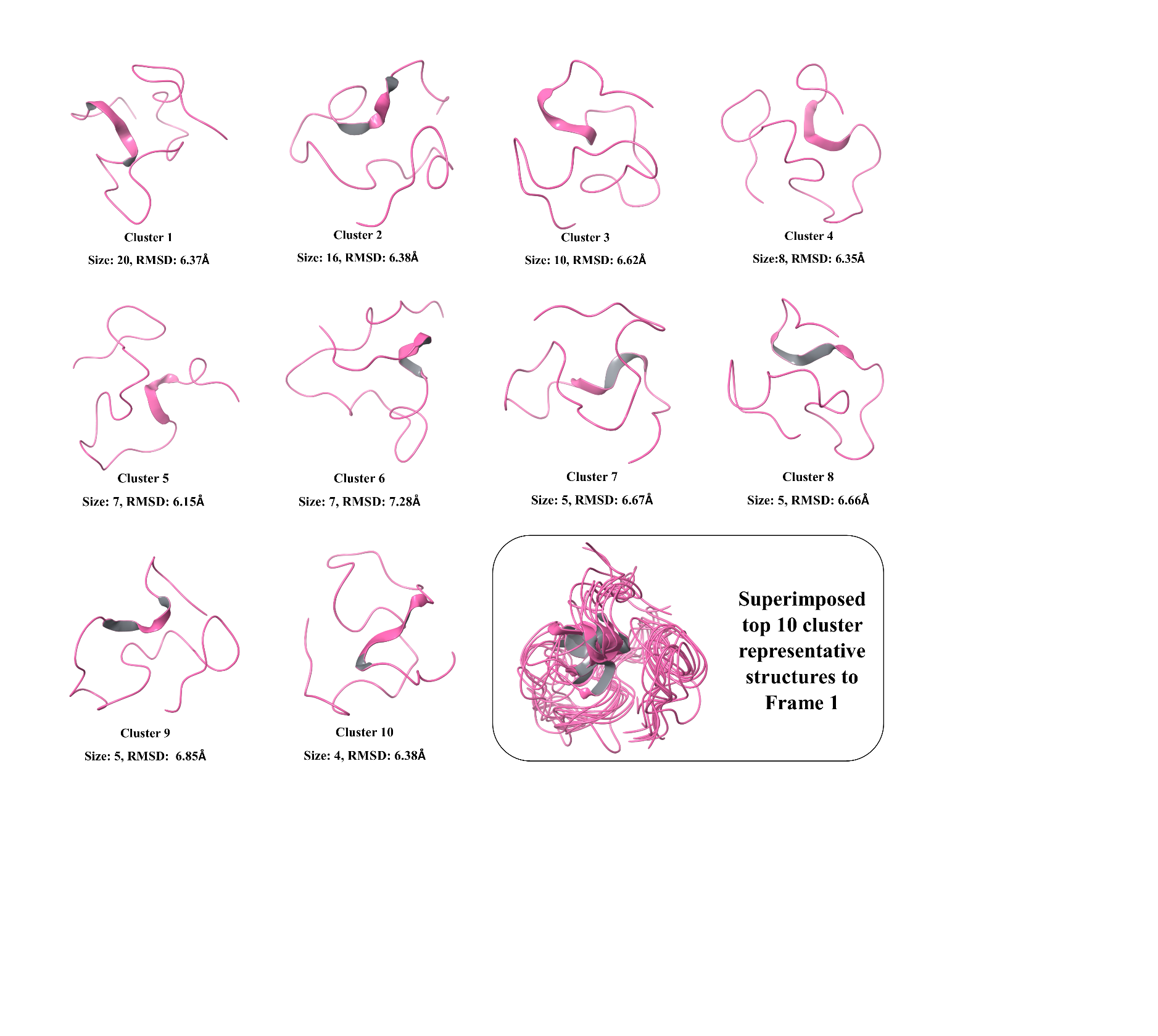

**Figure S10:** Representation of top 10 clusters from 1 µs simulation of NS4B-CR model B (I-TASSER) performed using OPLS2005 forcefield. Each cluster contains the certain frames (size) on the basis of RMSD calculated from the reference frame (frame 1). The superimposed structure shows all the cluster structured superimposed to frame 1.

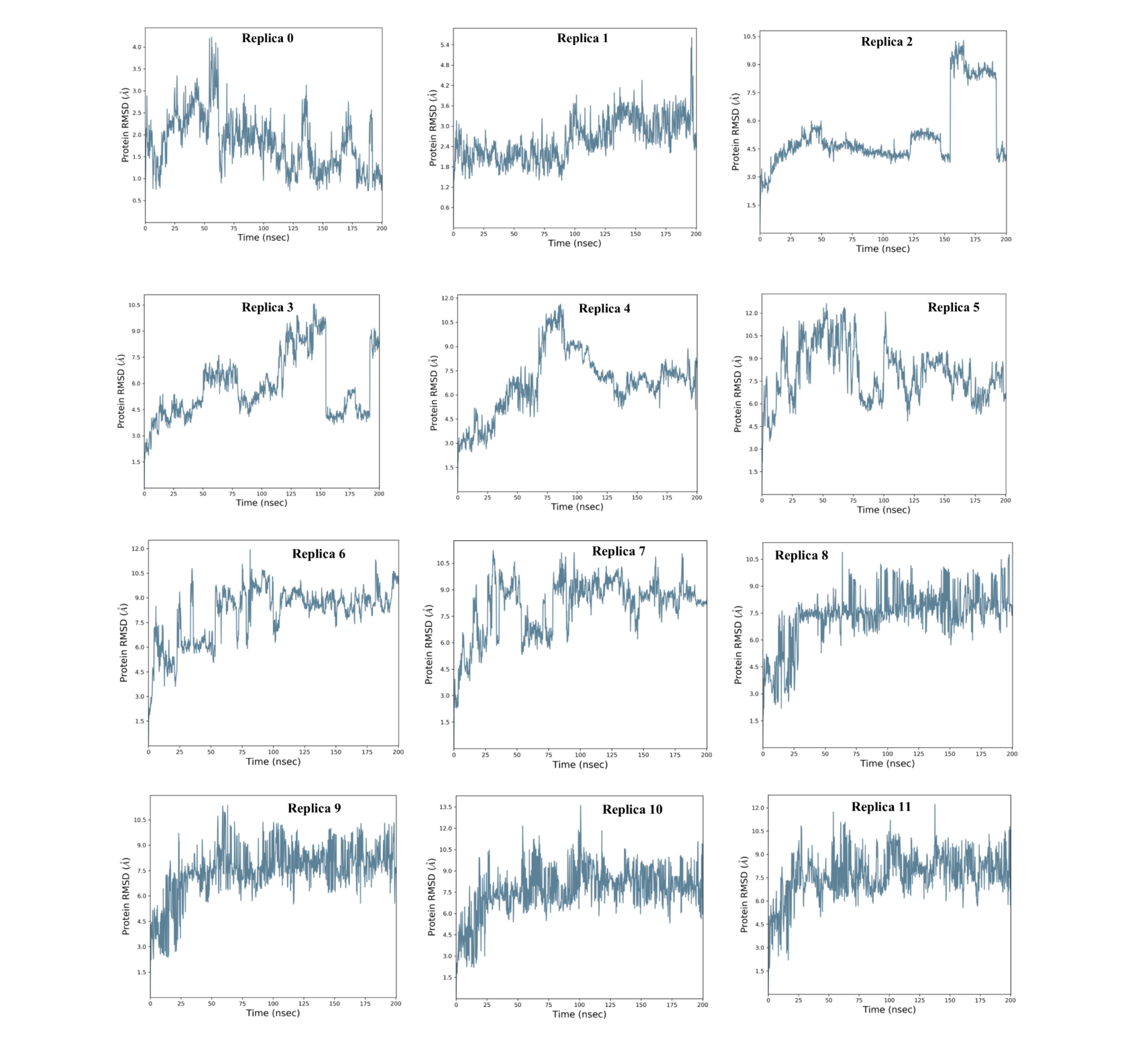

**Figure S11:** Graphs depicting the RMSD of replicas 0-11 from REMD simulations of simulated frame of model B (from I-TASSER simulation trajectory) of NS4B-CR.

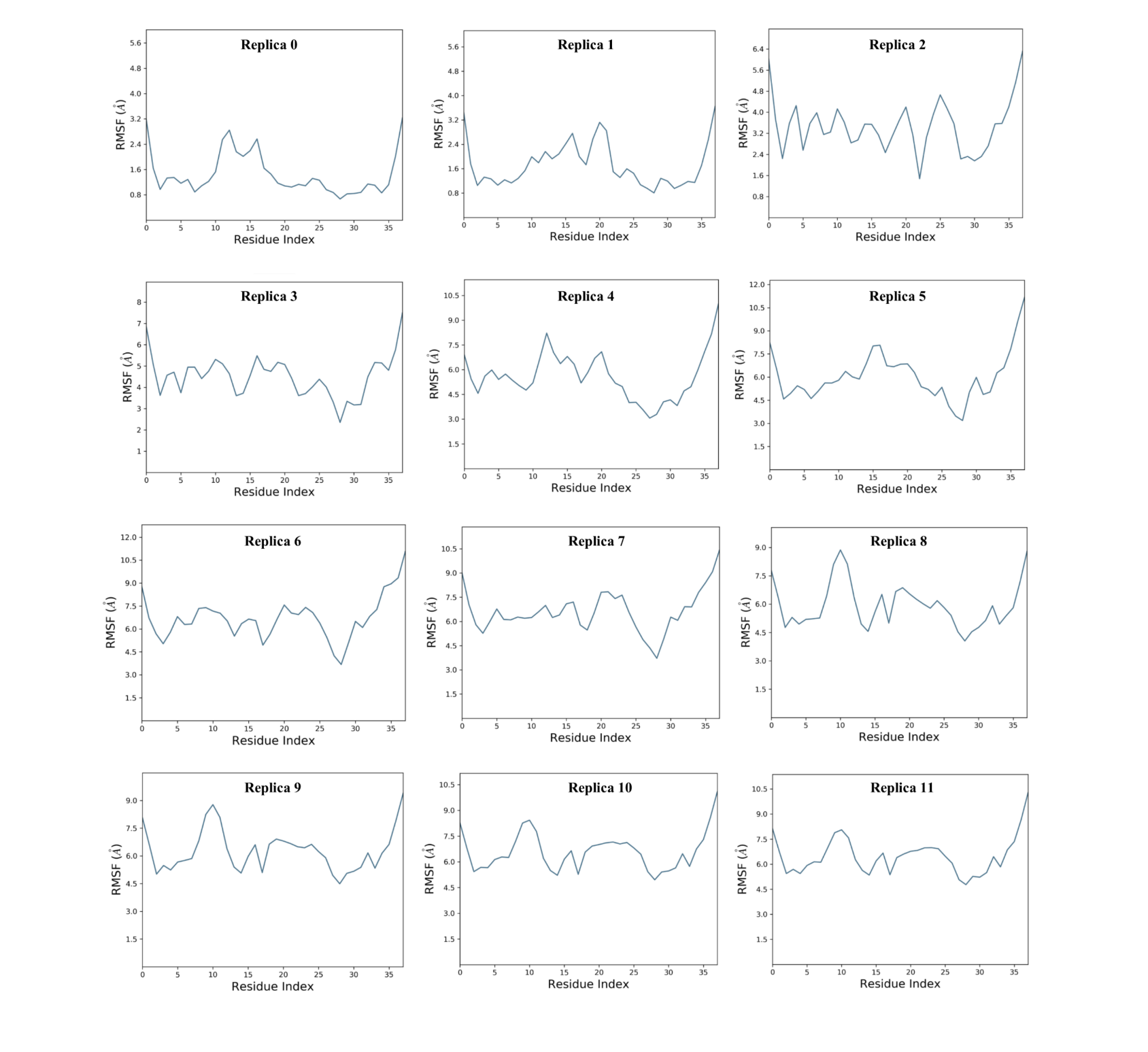

**Figure S12:** Graphs depicting the RMSF of replicas 0-11 from REMD simulations of simulated frame of model B (from I-TASSER simulation trajectory) of NS4B-CR.

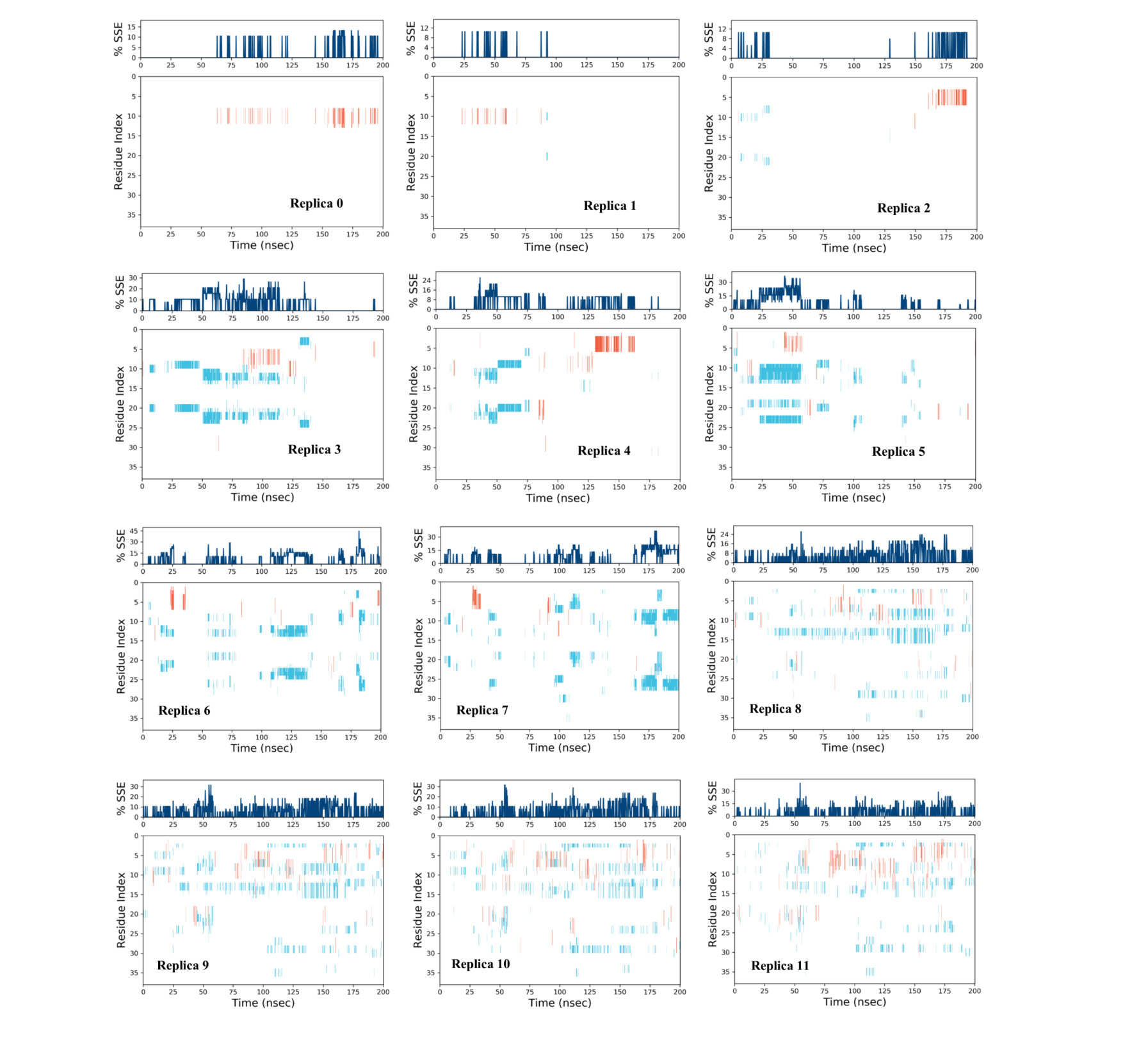

**Figure S13:** Secondary structure timelines of replicas 0-11 from REMD simulations of simulated frame of model B (from I-TASSER simulation trajectory) of NS4B-CR.

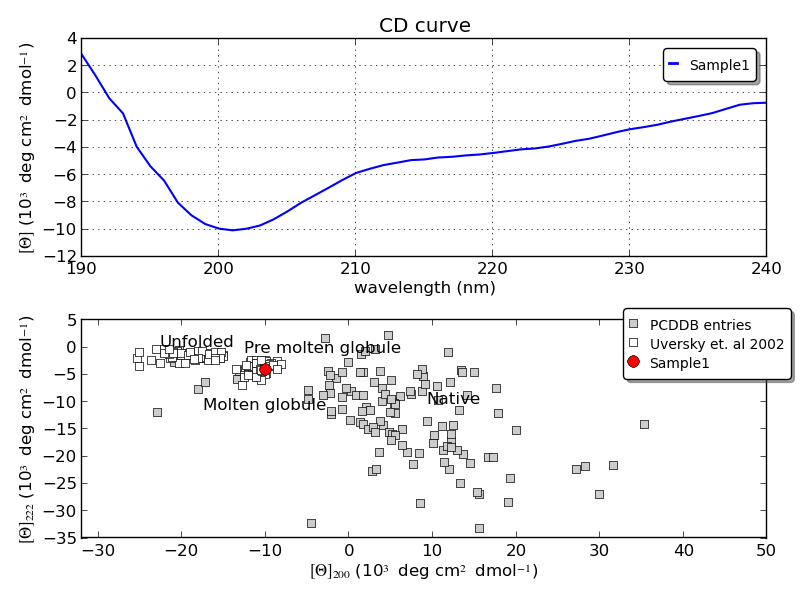

**Figure S14:** CAPITO analysis of CD spectra of NS4B-CR peptide in presence of 500mM SDS micelles. CD Spectra is shown in blue colour in above panel while negative ellipticity at 222 nm is depicted with red colour (sample 1).

**Supplementary Table**

**Table 1:** Secondary structure information present in each replica of NS4B-CR peptide in REMD simulation.

| **Replica** | **Temperature (K)** | **% α-Helix** | **% β-sheet** | **% Total SSE** |
| --- | --- | --- | --- | --- |
| **0** | 300 | 0.86 | 0 | 0.86 |
| **1** | 310 | 0.25 | 0.03 | 0.28 |
| **2** | 320 | 0.62 | 0.18 | 0.8 |
| **3** | 330 | 0.53 | 4.2 | 4.74 |
| **4** | 340 | 1.26 | 1.97 | 3.24 |
| **5** | 350 | 0.45 | 4.21 | 4.67 |
| **6** | 360 | 0.46 | 3.78 | 4.24 |
| **7** | 370 | 0.43 | 4.41 | 4.83 |
| **8** | 380 | 0.56 | 3.53 | 4.09 |
| **9** | 390 | 0.8 | 3.6 | 4.39 |
| **10** | 400 | 0.96 | 3.11 | 4.07 |
| **11** | 410 | 1.21 | 2.24 | 3.45 |

**Supplementary Movies**

**Movie 1:** Simulation trajectory (1 μs) of full-length ZIKV NS4B protein AlphaFold2 predicted model A in presence of POPC membrane.

**Movie 2:** Simulation trajectory (1 μs) of full-length ZIKV NS4B protein I-TASSER predicted model B in presence of POPC membrane.

**Movie 3:** Simulation trajectory (1 μs) of N-terminal of ZIKV NS4B AlphaFold2 predicted structure.

**Movie 4:** Simulation trajectory (1 μs) of C-terminal of ZIKV NS4B AlphaFold2 predicted structure in absence of POPC membrane.

**Movie 5:** Simulation trajectory (1 μs) of C-terminal of ZIKV NS4B AlphaFold2 predicted structure in presence of POPC membrane.

**Movie 6:** Simulation trajectory (1 μs) of Cytosolic region of ZIKV NS4B AlphaFold2 predicted model A structure.

**Movie 7:** Simulation trajectory (1 μs) of Cytosolic region of ZIKV NS4B I-TASSER predicted model B structure.
